# A dissection of the genomic antimicrobial resistance epidemiology of *Salmonella* Typhimurium

**DOI:** 10.1101/2024.05.12.593721

**Authors:** Sandeep Kaur, Michael Payne, Sally R. Partridge, Vitali Sintchenko, Ruiting Lan

## Abstract

*Salmonella* Typhimurium (STm) is a globally prevalent pathogen. We compiled a dataset comprising ∼65,000 publicly available STm isolates and analysed the predicted-resistance to 15 key antibiotics. Additionally, we typed all isolates using a standardized typing method, multilevel genome typing (MGT), and characterised the resistance profiles by MGT sequence types (ST). We identified 407 MGT STs wherein at least 80% of the isolates were non-susceptible to at least one antibiotic. For the key antibiotics prescribed for severe salmonellosis, we identified three ciprofloxacin non-susceptible MGT STs, and eight cefotaxime non-susceptible MGT STs in the last two years of the dataset (2021-2022). While the ciprofloxacin non-susceptible MGT STs comprised isolates predominantly from the UK; only one cefotaxime non-susceptible MGT ST comprised isolates predominantly from UK and associated with swine, with others from USA, and associated with cattle and poultry. Integration of AMR predictions with MGT strain typing provides sharable, standardised, and specific identification and tracking of resistant isolates. This integrated analysis presents a unique approach for the global surveillance of antimicrobial resistance and resistant strains.

## Introduction

Globally, *Salmonella enterica* serovar Typhimurium (STm) is one of the two most frequently isolated non-typhoidal *Salmonellae* (NTS) serovars [1, 2]. NTS, especially STm, has a broad host range and infects humans through direct contact with hosts or consumption of contaminated food [3, 4]. NTS infects the gastrointestinal tract, and disease severity is generally mild to moderate. However, NTS infections can also result in bacteraemia, and cause endocarditis, abscesses, or meningitis [5]. Antibiotics are not recommended for mild to moderate cases of salmonellosis in healthy individuals but are required in severe cases. For such cases, the US CDC recommends fluoroquinolones (e.g., ciprofloxacin) as first-line treatment in adult travellers [6]. Ceftriaxone (third-generation cephalosporin) or azithromycin (macrolide) are recommended for children or adults with invasive disease.

By multilocus sequence typing (MLST), STm is dominated by four sequence types (STs), ST19, ST34, ST313 and ST36 [7]. ST19 predominates, while ST34 contains the monophasic variant of STm. ST313 includes two multidrug resistant lineages (L1 and L2) associated with invasive disease in Africa [8]. ST36 has been shown to form a distinct phylogenetic group from other STs within STm [9, 10].

In our previous work, we developed multilevel genome typing (MGT) for characterising STm isolates at multiple levels of resolution [11]. The MGT scheme consists of nine MLST-like schemes that increase in resolution from MGT1 to MGT9. Hence, a set of nine MGT STs, one for each level, are assigned to every isolate, and via these nine STs, isolates can be studied at multiple resolutions (such as global, local, long-term or short-term). MGT1 is the traditional 7-gene MLST for the *Salmonella* species and MGT1 STs correspond to the established 7-gene MLST STs, e.g., ST19 or ST34. MGT8 comprises the species core genome MLST (cgMLST). MGT2 to 7 are composed of increasing numbers of mutually-exclusive sets of core genes from MGT8. MGT9 is the serovar core genome MLST. Many well studied clades and types can be described using intermediate resolution levels. MGT3 ST11 describes the phage type DT104, a global pandemic clone, while two large groups composed mainly of avian isolates from the USA belong to MGT3 ST8 or MGT3 ST171. Monophasic STm within ST34, corresponds to MGT3 ST26, MGT3 ST18 or MGT3 ST225. The invasive disease-causing lineages within ST313 (L1 and L2) correspond to MGT3 ST725 and MGT3 ST3, respectively.

The emergence of antimicrobial resistance (AMR) and multi-drug resistance (MDR) in STm has been well documented [4, 12]. For example, ST34 isolates often have the ASSuT resistance pattern (i.e. to **a**mpicillin, **s**treptomycins, **su**lphathiazoles, and **t**etracyclines), with the corresponding genes *bla*_TEM-1_, *strA*-*strB* (or *aph(3’’)-Ib* and *aph(6)-Id*), *sul2*, and *tet*(B) present on a chromosomal resistance island [13]. ST34 isolates may also be resistant to additional drugs, such as quinolones, cefotaxime, colistin, gentamicin, and chloramphenicol [14]. The resistance pattern ACSSuT (the additional ‘C’ refers to **c**hloramphenicol) has been found in a large proportion of phage type DT104, or MGT3 ST11, isolates. The ACSSuT genes have been reported as *tet*(G), *tet*R, *aadA2*, *bla*_CARB-2_, *sul1*, *floR*, present within the chromosomal *Salmonella* genomic island 1 (SG1) [15–17]. In DT104, resistance to additional drugs such as gentamicin, kanamycin, trimethoprim-sulphamethoxazole has also been observed [18]. The DT104 clone was characterized as a pandemic strain from 1990 to 2010, whereas ST34 (or DT193/DT120) is characterised as the current pandemic strain [4]. The prevalent lineage associated with bloodstream infections, ST313 L2, or MGT3 ST3, has well known resistance to multiple drugs such as ampicillins, trimethoprim-sulphamethoxazole and chloramphenicol with reports also emerging of resistance to azithromycin and cephalosporins [8].

In recent years, whole genome sequencing (WGS) has been adopted for the surveillance of *Salmonella* in the UK [19] and the USA [20] with complete WGS surveillance datasets available publicly. Given this availability of a large number of genomes from long term *Salmonella* surveillance projects, we conducted a comprehensive survey of AMR in STm. We particularly focussed on combining detection of AMR determinants with MGT typing. We demonstrated the utility of the standardised and multi-resolution MGT approach to precisely identify groups of non-susceptible strains, especially those with resistance to clinically important drugs, and determined global temporal trends of AMR MGT STs in STm.

## Results

### Dataset

We previously developed and maintain a database (MGTdb) for typing STm isolates using MGT [11, 21]. Our dataset consisted of 65,859 genomes. It was composed of publicly available isolates within MGTdb, that were assigned a ST at at-least one MGT level, until August 31, 2023. In this dataset, over half of the isolates had metadata (**Table 1**), 63% with country, 59% with year, and 58% with both country and year. The isolates spanned 94 countries and were sampled between 1900 and 2022. Since, around 2013, the number of available whole-genome sequenced isolates sharply increased (**Figure S1**). Of the available isolates with both year and country, 73% were from the USA or the UK.

**Table 1:** Number of isolates from top 10 countries in descending order.

| Country | With country |  | With country and year |  | 2015-2018 |  | 2019-2022 |  |
| --- | --- | --- | --- | --- | --- | --- | --- | --- |
|  | Count | % | Count | % | Count | % | Count | % |
| USA | 15643 | 37.9 | 13519 | 35.1 | 4791 | 32.5 | 5216 | 42.2 |
| United Kingdom | 15117 | 36.6 | 14733 | 38.3 | 7127 | 48.4 | 5066 | 41 |
| Canada | 1737 | 4.2 | 1692 | 4.4 | 840 | 5.7 | 130 | 1.1 |
| Australia | 1557 | 3.8 | 1557 | 4 | 494 | 3.4 | 121 | 1 |
| Denmark | 1148 | 2.8 | 1141 | 3 | 31 | 0.2 | 4 | 0 |
| South Africa | 1049 | 2.5 | 1049 | 2.7 | 32 | 0.2 | 992 | 8 |
| Germany | 959 | 2.3 | 959 | 2.5 | 544 | 3.7 | 96 | 0.8 |
| Taiwan | 483 | 1.2 | 483 | 1.3 | 115 | 0.8 | 1 | 0 |
| Italy | 332 | 0.8 | 332 | 0.9 | 88 | 0.6 | 3 | 0 |
| Japan | 319 | 0.8 | 319 | 0.8 | 136 | 0.9 | 0 | 0 |
| **Total displayed** | 38344 | 92.9 | 35784 | 93 | 14198 | 96.4 | 11629 | 94.1 |
| **Total overall** | 41285 |  | 38482 |  | 14738 |  | 12361 |  |
| **%** | 62.7 |  | 58.4 |  | 22.4 |  | 18.8 |  |

Since, the UK adopted genomic surveillance for *Salmonella* from 2015 [19], and the USA from 2019 [20], to provide a comparison between USA and UK that is potentially less biased by sampling effects, we considered a data subset ranging between 2019 to 2022, and three locations UK, USA and other countries (designated as ‘Other’). Additionally, as UK surveillance began from 2015, the 2015 to 2018 data could be compared within UK, hence, we considered another subset between 2015 to 2018, and similarly the three locations. Note that for the Other data, in both time-frames considered, since the temporal and geographic coverage of the isolates was very heterogenous, we present the results but refrain from discussing their significance, as any trends observed may be the result of sampling bias. This is similarly the case for USA between 2015 and 2018, however, as a large number of isolates are available from this country, we compare trends within the country itself over the two timeframes.

### Non-susceptibility to any antibiotic

A non-susceptibility score (NS_s_) for each isolate was defined as the number of antibiotic classes an isolate was non-susceptible to, out of the15 examined. Approximately half (47%, avg. NS_s_=2, avg. NS_s_ of non-susceptible isolates only=4.2) of the isolates were non-susceptible to at least one drug (**Table 2**), however this varied by both location and time (**Table 2**).

**Table 2:** Non-susceptibility score in the complete dataset, and in subsets of isolates from three locations in two timeframes.

| Non-susceptibility score | Overall |  | UK |  |  |  | USA |  |  |  | Other |  |  |  |
| --- | --- | --- | --- | --- | --- | --- | --- | --- | --- | --- | --- | --- | --- | --- |
|  |  |  | 2015-2018 |  | 2019-2022 |  | 2015-2018 |  | 2019-2022 |  | 2015-2018 |  | 2019-2022 |  |
|  | # | % | # | % | # | % | # | % | # | % | # | % | # | % |
| 0 | 35013 | 53.1 | 2600 | 36.5 | 2296 | 45.3 | 2777 | 58.0 | 2734 | 52.4 | 1491 | 50.4 | 1582 | 66.9 |
| 1 | 3274 | 5 | 638 | 9 | 337 | 6.7 | 134 | 2.8 | 150 | 2.9 | 189 | 6.4 | 141 | 6.0 |
| 2 | 3852 | 5.8 | 188 | 2.6 | 160 | 3.2 | 804 | 16.8 | 979 | 18.8 | 142 | 4.8 | 46 | 1.9 |
| 3 | 2412 | 3.7 | 482 | 6.8 | 331 | 6.5 | 104 | 2.2 | 95 | 1.8 | 141 | 4.8 | 181 | 7.7 |
| 4 | 10568 | 16 | 2065 | 29 | 1182 | 23.3 | 560 | 11.7 | 868 | 16.6 | 449 | 15.2 | 148 | 6.3 |
| 5 | 3133 | 4.8 | 379 | 5.3 | 287 | 5.7 | 171 | 3.6 | 152 | 2.9 | 172 | 5.8 | 81 | 3.4 |
| 6 | 3981 | 6 | 297 | 4.2 | 131 | 2.6 | 140 | 2.9 | 116 | 2.2 | 118 | 4.0 | 50 | 2.1 |
| 7 | 1593 | 2.4 | 225 | 3.2 | 113 | 2.2 | 36 | 0.8 | 47 | 0.9 | 74 | 2.5 | 36 | 1.5 |
| 8 | 957 | 1.5 | 157 | 2.2 | 75 | 1.5 | 28 | 0.6 | 20 | 0.4 | 85 | 2.9 | 54 | 2.3 |
| 9 | 638 | 1 | 53 | 0.7 | 133 | 2.6 | 12 | 0.3 | 21 | 0.4 | 46 | 1.6 | 35 | 1.5 |
| 10 | 360 | 0.5 | 31 | 0.4 | 18 | 0.4 | 16 | 0.3 | 23 | 0.4 | 37 | 1.3 | 5 | 0.2 |
| 11 | 75 | 0.1 | 8 | 0.1 | 2 | 0.0 | 5 | 0.1 | 11 | 0.2 | 12 | 0.4 | 2 | 0.1 |
| 12 | 39 | 0.1 | 4 | 0.1 | 1 | 0.0 | 4 | 0.1 | 0 | 0.0 | 3 | 0.1 | 3 | 0.1 |
| 13 | 0 | 0 | 0 | 0 | 0 | 0.0 | 0 | 0.0 | 0 | 0.0 | 0 | 0.0 | 0 | 0.0 |
| 14 | 0 | 0 | 0 | 0 | 0 | 0.0 | 0 | 0.0 | 0 | 0.0 | 0 | 0.0 | 0 | 0.0 |
| 15 | 0 | 0 | 0 | 0 | 0 | 0.0 | 0 | 0.0 | 0 | 0.0 | 0 | 0.0 | 0 | 0.0 |

In the three location datasets (UK, USA and Other) from 2015-2022, the isolates from UK had the highest non-susceptibility, with 59% of isolates non-susceptible to at least one drug (avg. NS_s_=2.4, avg. NS_s_ of non-susceptible isolates only=4), 45% of USA isolates were non-susceptible to at least one drug (avg. NS_s_=1.5, avg. NS_s_ of non-susceptible isolates only=3.4), and 41% of Other isolates were non-susceptible to at least one drug (avg. NS_s_=1.8, avg. NS_s_ of non-susceptible isolates only= 4.3). Additional results comparing non-susceptibility are detailed in **Section Supplementary Results**.

### Non-susceptibility to individual antibiotics

Non-susceptibility to any individual antibiotic in the complete dataset varied from negligible, with 0.02% of isolates predicted to be resistant to meropenem, to 36% of isolates predicted to be resistant to tetracycline (**Figure 1)**. The UK in 2015-2018 had some of the highest proportion of resistant isolates, especially for tetracycline (54%), streptomycin (52%), ampicillin (49%) and sulfathiazole (46%). In the UK there was a significant increase in the proportion of resistant isolates for only two antibiotics, kanamycin (from 2.9% to 5%) and gentamicin (from 1.7% to 2.9%) in 2019-2022 compared to 2015-2018, and a significant reduction in the proportion of resistant isolates for eight antibiotics namely, tetracycline, streptomycin, ampicillin, sulfathiazole, cefotaxime, azithromycin, colistin, and aminoglycosides (Rmt) (**Figure 1**). In the USA, the proportion of resistant isolates for four antibiotics increased significantly in 2019-2022 compared to 2015-2018, and decreased for three antibiotics. In particular, non-susceptibility to tetracycline (38% to 44%), streptomycin (19% to 22%), sulphathiazole (33% to 39%) and gentamicin (4.3% to 8.2%) increased; and to chloramphenicol, cefotaxime, and aminoglycosides (Rmt), decreased. Although the proportion of isolates with resistance to ciprofloxacin remained unchanged in both the UK and the USA, the proportion of intermediately-resistant isolates significantly decreased in the UK (9% to 8%), and increased in the USA (from 2% to 3%). Additional results comparing non-susceptibility to individual antibiotics in the UK and the USA are detailed in **Supplementary Results.**

**Figure 1.**
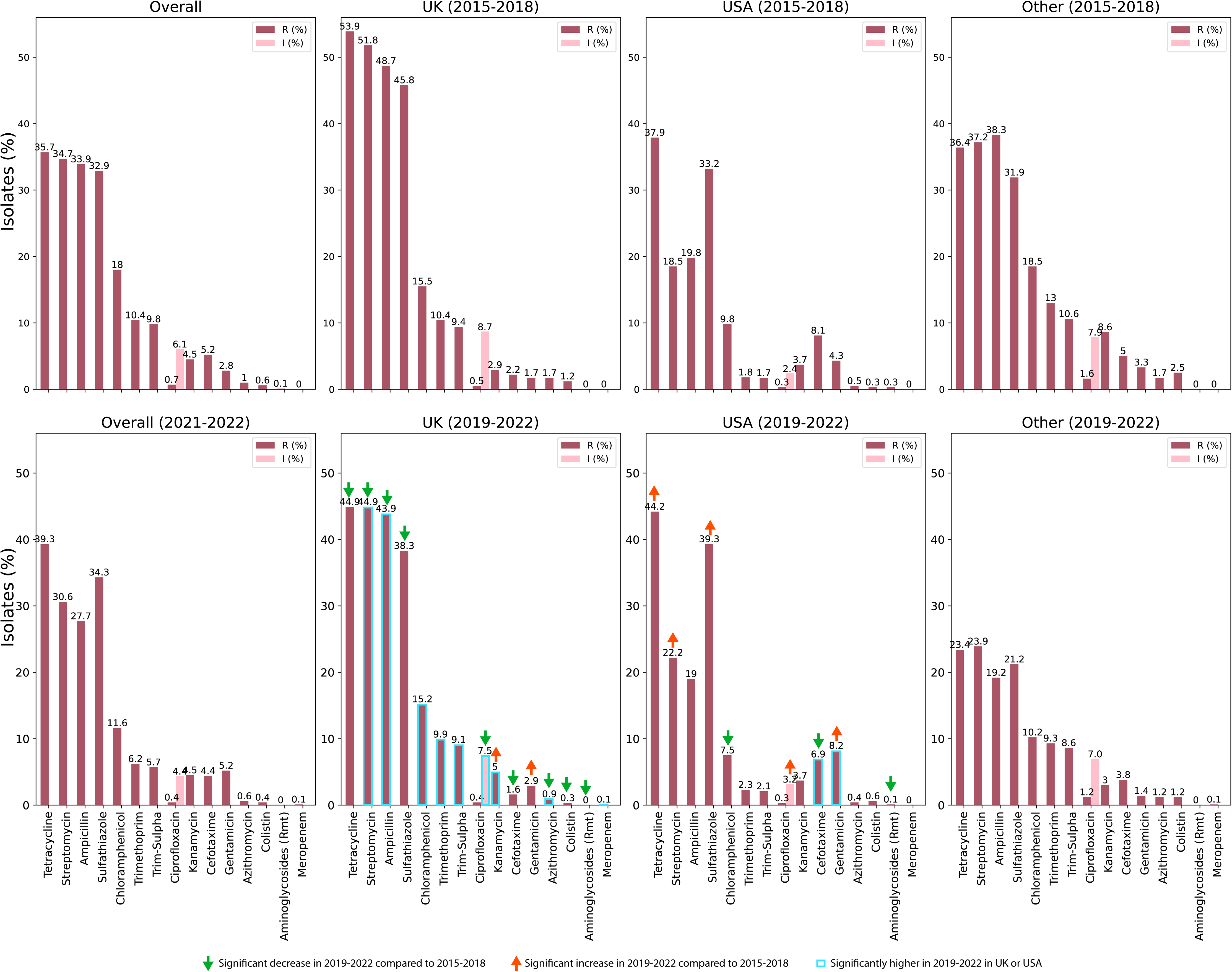
Non-susceptibility to 15 antibiotics within the complete dataset and various subsets. In each plot, the antibiotics have been arranged from left to right in descending order of non-susceptibility in the complete dataset. Significantly higher non-susceptibility in 2019-2022, compared to 2015-2018, in the isolates from UK and USA, is indicated via an upwards red arrow, and significantly lower non-susceptibility is indicated via a downwards green arrow. Significantly higher non-susceptibility between UK and USA in 2019-2022 is indicated via a blue outline around the bar. Non-susceptibility to an antibiotic between two groups was compared using the Binomial test. Significance was assessed at the Bonferroni corrected *p* < 0.0008.

### Non-susceptibility within MGT1 (MLST) STs

MGT1 (MLST) ST19 was the most frequent ST (71% isolates) in the dataset followed by ST34 (14%), then ST313 (5%) and ST36 (3%) (**Figure 2**, **Table S1**). As expected, most isolates in ST19, were susceptible to all drugs (64%), whereas most isolates in ST34 were non-susceptible to atleast one drug (96%). Similarly, ST313 isolates were predominantly non-susceptible (81%), whereas ST36 isolates were predominantly susceptible (73%). Comparison of the average non-susceptibility score of isolates within these STs revealed the following order of resistance: ST313 (avg. NS_s_=4.8) > ST34 (avg. NS_s_=4) > other STs (avg. NS_s_=1.7) > ST19 (avg. NS_s_=1.4) > ST36 (avg. NS_s_=1.3). When comparing the average non-susceptibility score of non-susceptible isolates only, ST313 (avg. NS_s_ of non-susceptible isolates only=6) had the highest non-susceptibility, followed by ST36 (avg. NS_s_ of non-susceptible isolates only=4.8) and ST34 (avg. NS_s_ of non-susceptible isolates only=4.1), then by other STs (avg. NS_s_ of non-susceptible isolates only=4.2) and ST19 (avg. NS_s_ of non-susceptible isolates only=3.9). The differences between ST36 and ST34, and other STs and ST19 were assessed as not significantly different.

**Figure 2.**
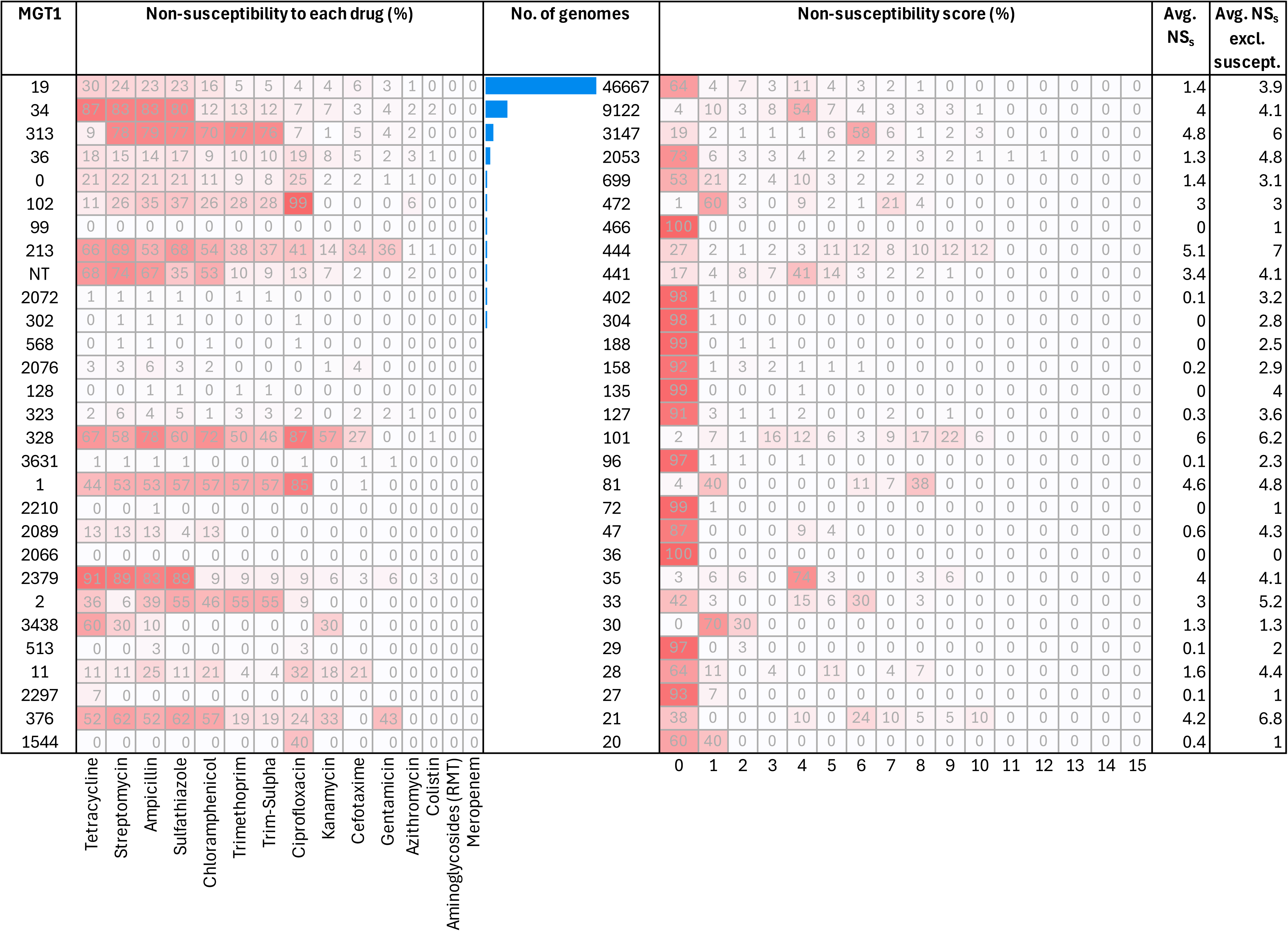
Non-susceptibility to 15 antibiotics and other aggregate non-susceptibility metrics for MGT1 STs. The MGT1 STs are ordered in descending order of number of genomes – and only STs with a minimum count of 20 isolates are shown. ‘0’ indicates a new ST, NT (or Not Typable) indicates that an ST was not assigned (for example due to missing data). Non-susceptibility score for each isolate is the count of antibiotics for which an isolate was non-susceptible. Thus, the heatmap ‘Non-susceptibility score (%)’ depicts the distribution of the non-susceptibility (NS) score across the 15 antibiotics for each MGT1 ST. ‘Avg. NS_s_’ indicates the average non-susceptibility score of all isolates within the MGT1 ST, and ‘Avg. NS_s_ excl. suscept.’ indicates the average non-susceptible score of isolates with non-susceptibility to at least one antibiotic, i.e. excluding isolates which were identified as susceptible to all 15 antibiotics.

Additional results comparing non-susceptibility to individual antibiotics within MGT1 STs are detailed in **Supplementary Results**. Additionally, results of the distribution of the ASSuT phenotypic and genotypic resistance pattern in MGT1 ST34, and the distribution of the ACSSuT phenotypic and genotypic resistance pattern in MGT3 ST11 (or DT104) are also presented in **Supplementary Results**.

### Highly non-susceptible MGT STs to a given antibiotic

Analysis of AMR using MGT1 STs gave a broad overview of non-susceptibility in the dataset, especially in the context of the most frequent STs, ST19 and ST34. However, to obtain a finer-grade understanding of relationships between genotypes and AMR, we used MGT to classify isolates into higher-resolution MGT STs. We then identified a set of MGT STs, comprising sets of mutually-exclusive isolates, with very high non-susceptibility. In particular, we defined a “non-susceptible MGT ST” for an antibiotic, as a MGT ST wherein atleast 80% of the isolates were resistant to that antibiotic (see **Supplementary Results** for details on why the 80% cutoff was chosen).

In the complete dataset, we identified 407 non-susceptible MGT STs (**Table S2**, **Table 3, Supplementary File 1**). Non-susceptible MGT STs were identified for all antibiotics except aminoglycoside (Rmt) and meropenem, and spanned all MGT levels, from MGT1 to MGT9 (**Table 3, Figure S8)**. The majority of the MGT STs were non-susceptible to 1 drug (146 of 407), with 1 MGT ST non-susceptible to 10 drugs (**Table S2**).

**Table 3.**
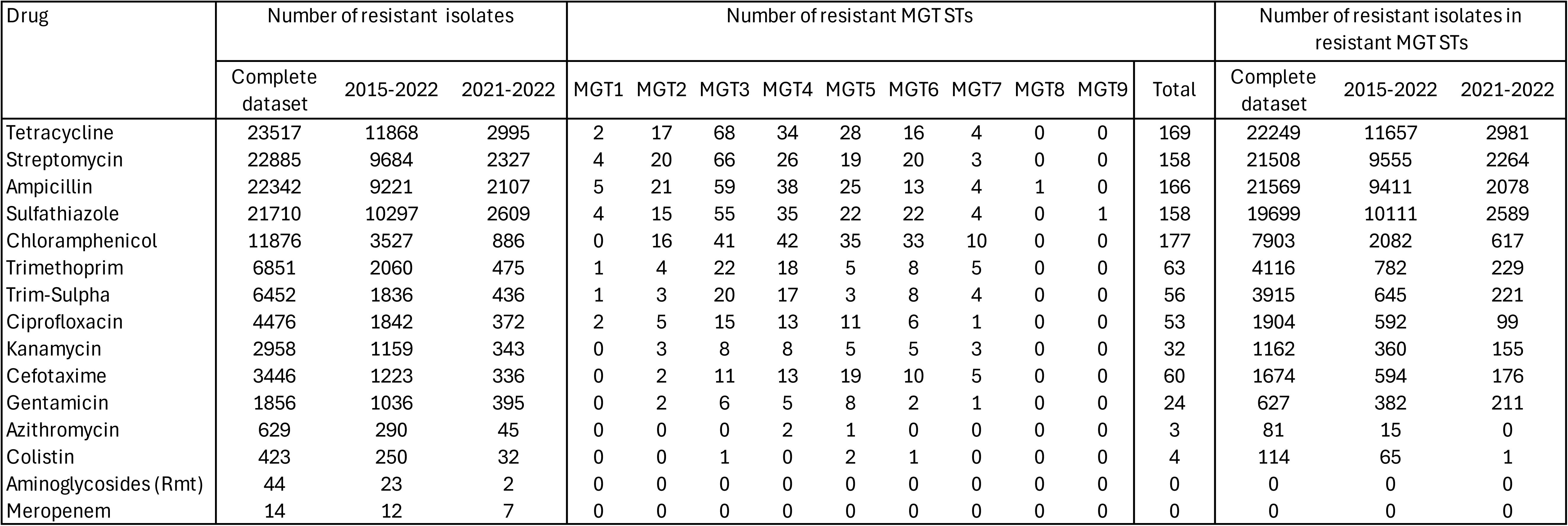
Number of resistant MGT STs identified at each MGT level for each antibiotic.

### Resistance to prescribed and last-line antibiotics

Non-susceptible MGT STs for commonly prescribed antibiotics (namely, ciprofloxacin, cefotaxime, and azithromycin), and last-line antibiotics (namely, carbapenems and colistin), are discussed in this paper. In particular, for both ciprofloxacin and cefotaxime, we identified major STs, i.e STs with minimally 10 isolates, in 2021-2022 (last two years in the dataset). Thus, non-susceptible MGT STs for these two drugs are detailed below, with a particular focus on non-susceptibility in 2021-2022. For colistin and azithromycin, major MGT STs were not present in 2021 to 2022; and for meropenem, only 14 isolates in the entire dataset were identified as resistant; and hence the details for resistance to these antibiotics are presented in the **Supplementary Results**. Lastly, we also assessed the reliability of non-susceptibility prediction, especially for cefotaxime and ciprofloxacin, and these results are also detailed in **Supplementary Results**.

### Ciprofloxacin

We identified 53 ciprofloxacin non-susceptible MGT STs which comprised 1,840 of 4,476 (41%) of ciprofloxacin non-susceptible isolates (**Figure S8**), and included 64 susceptible isolates. In these 53 MGT STs, 906 isolates (48%) contained a year annotation, using which 20 MGT STs were identified as major STs in 2015 to 2022. The largest ST (MGT2 ST59) comprised 120 isolates within this timeframe, and the remaining STs averaged around 18 isolates per ST.

We temporally-clustered these 20 MGT STs to identify the temporal patterns within this set. We identified seven distinct temporal patterns within the 20 MGT STs (**Figure 3a**). To enable comparison between the STs, the MGT STs were standardized around the mean count of isolates per year. Thus, each observed increase or decrease for an ST represents fold change around the mean. For example, the number of isolates of MGT2 ST59 (Cluster 4) increased between 2016 and 2017 (by approximately 1.5 fold), then dropped below average in 2020 to 2021, and was making a resurgence in 2022 (where the isolate counts are approximately the average for that ST). The clusters were temporally ordered to represent when the STs within the cluster first peaked – for example Clusters 1 and 2 contain MGT STs that were already present in 2015, whereas Cluster 7 represents MGT STs that arose in 2021. Four clusters comprised isolates mostly from UK (namely Clusters 1,3,4 and 7), two clusters comprised isolates mostly from UK and Other (Clusters 2 and 5, with isolates from China, Mexico, Colombia, USA and Australia), and Cluster 6 comprised isolates only from Other (in particular, from China).

**Figure 3.**
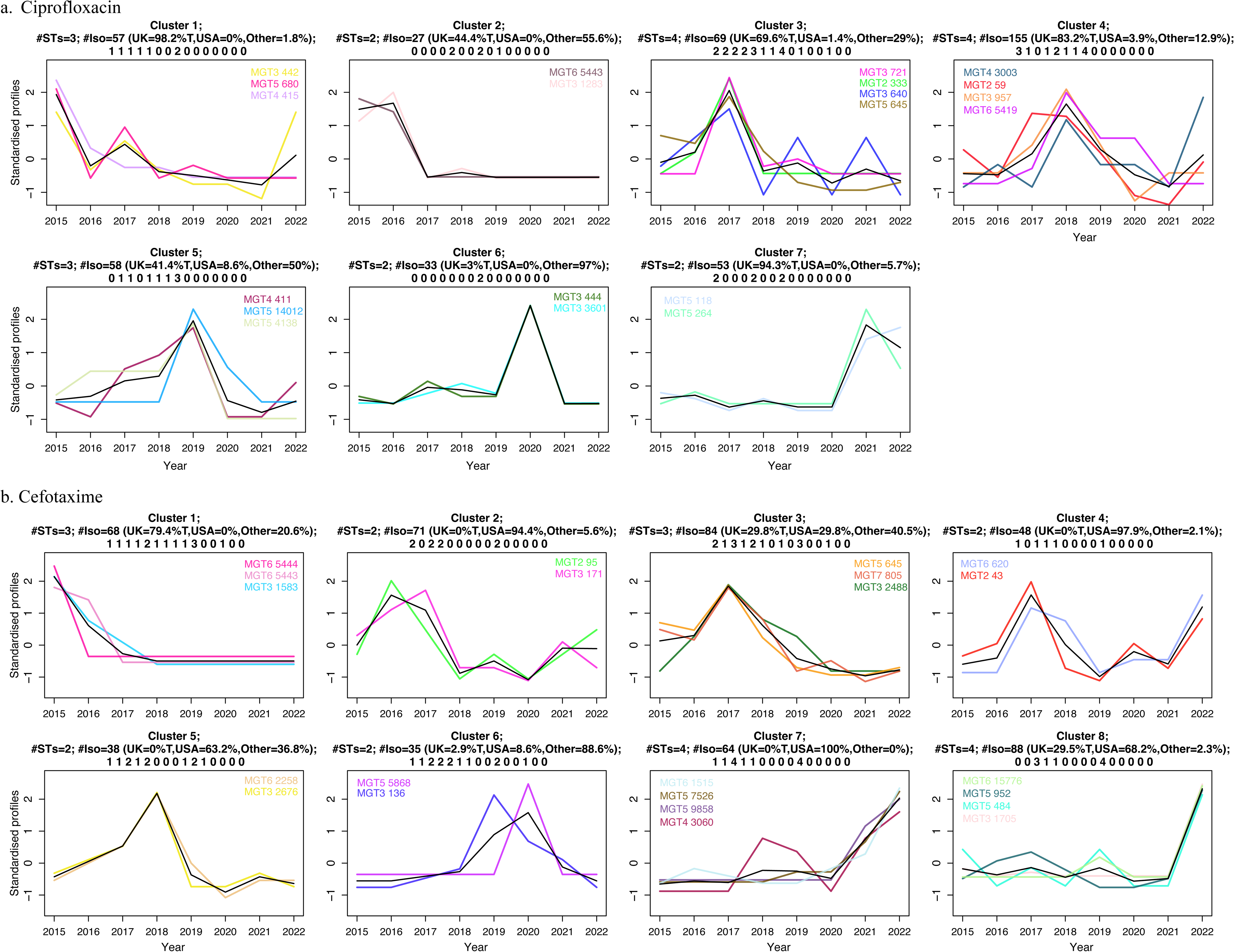
Temporal trends between 2015-2022 of non-susceptible MGT STs. a. Shows 20 ciprofloxacin non-susceptible MGT STs grouped into seven clusters, and b. shows 22 cefotaxime non-susceptible MGT STs grouped into eight clusters. Each cluster represents a unique temporal pattern, indicated by the bold-black line, which represents a weighted average of all MGT STs within the cluster. Each uniquely colored line represents the temporal profile of an MGT ST. The temporal profile of an MGT ST is the standardized count of isolates per year. The clusters have been ordered according to when the cluster first peaked in time. The cluster header indicates the number of MGT STs, the total number of isolates, and the geographical distribution of the isolates within the cluster. This is followed by counts in the order: “tetracycline, streptomycin, ampicillin, sulphathiazole, chloramphenicol, trimethoprim, trimethoprim-sulphathiazole, ciprofloxacin, kanamycin, cefotaxime, gentamicin, azithromycin, colistin, aminoglycosides (Rmt) and meropenem”, representing the number of MGT STs identified as non-susceptible to the antibiotic at that position within the cluster. The clusters were generated using the unsupervised *c*-means clustering algorithm, and the optimal number of clusters were identified using both a combination of the silhouette index and manual inspection of the temporal-delineation of the clusters.

In 2021-2022, last two years with year annotation available, only three MGT STs comprised at least 10 isolates, namely MGT2 ST59, MGT5 ST118 and MGT5 ST264. While all three MGT STs were isolated predominantly from UK, MGT2 ST59 was also isolated in the USA and other countries including India, Denmark, Canada, and Australia. Both the MGT5 STs were also found in other European countries (such as Denmark). All three of these MGT STs were within MGT1 ST19, and the MGT5 STs were within MGT3 ST11 or DT104 (**Figure 4a**) - with MGT5 ST118 having a 13% genotypic pattern of resistance of the well known ACSSuT genes, and similarly for MGT5 ST264 at 33% for. Although the host range for most isolates were missing, all three STs had isolates with annotations as isolated from human. Additionally, both MGT5 STs were also isolated from bovine sources (**Figure 4a**).

**Figure 4.**
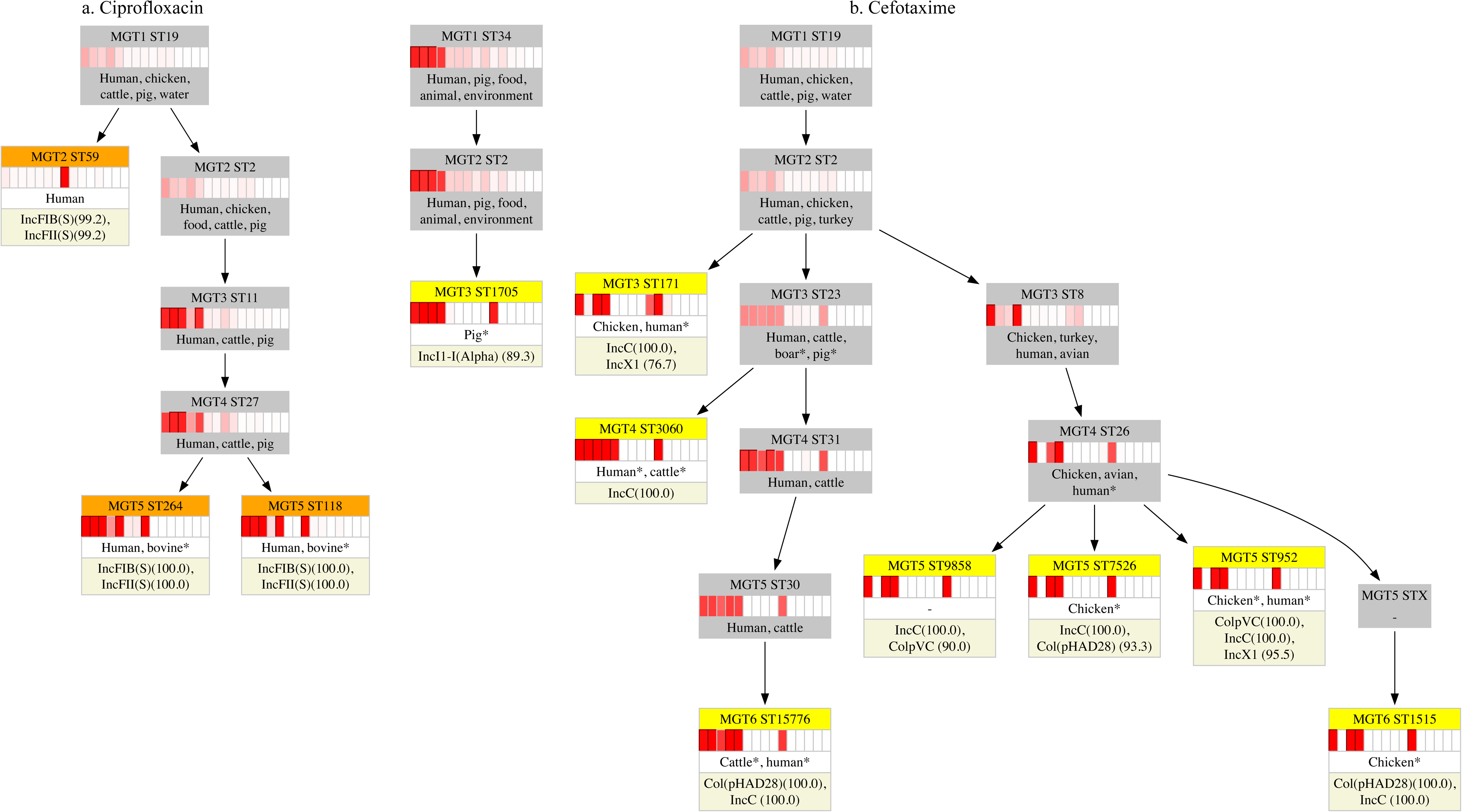
MGT hierarchy of non-susceptible MGT STs with at least 10 isolates in 2021-2022. a. Shows three ciprofloxacin non-susceptible MGT STs (in orange), and b. shows eight cefotaxime non-susceptible MGT STs (in yellow). For any MGT ST level 2 or higher, the lower level MGT ST is displayed when at least 50% isolates in the current MGT level ST are assigned the same ST at the lower MGT level, otherwise an ‘X’ is displayed. The heatmap indicates the percentage of isolates within the ST that are non-susceptible to antibiotics in the order: “tetracycline, streptomycin, ampicillin, sulphathiazole, chloramphenicol, trimethoprim, trimethoprim-sulphathiazole, ciprofloxacin, kanamycin, cefotaxime, gentamicin, azithromycin, colistin, aminoglycosides (Rmt) and meropenem”. A black outline is indicated for an antibiotic in the heatmap when at least 80% of the isolates were non-susceptible to the antibiotic at that position. Below the heatmap, the most frequent (top five, where available) isolation sources of the isolates in the ST are indicated, with an asterisk indicating that less than 10 isolates were annotated with this source. The sources were manually modified to increase clarity. Following the source, for the non-susceptible MGT STs, the last row shows the plasmid replicon types, followed by the percentage of isolates within which the replicon types were identified. Only those replicon types are shown that were present in at least 50% of the isolates in the ST.

With regards to the underlying mechanism of resistance, of these 53 MGT STs, there were 14 combinations of mutations or genes (**Figure S9**). 18 MGT STs had some level of complete resistance, ranging from 100% in MGT2 ST333 and 4% in MGT1 ST1, and the remaining 35 were only intermediately resistant (or require increased exposure [22]) to ciprofloxacin. In particular, for the MGT STs isolated in 2021 to 2022 (**Figure 5a**): MGT5 ST118 and MGT5 ST364 were only intermediately resistant to ciprofloxacin via the mutation *gyrA* D87N. MGT2 ST59 was also predominantly intermediately resistant with all non-susceptible isolates containing the mutation *gyrA* D87Y, however 4% isolates were ciprofloxacin resistant, and in addition to the mutation contained the genes *qnrS13* or *qnrS1*.

**Figure 5.**
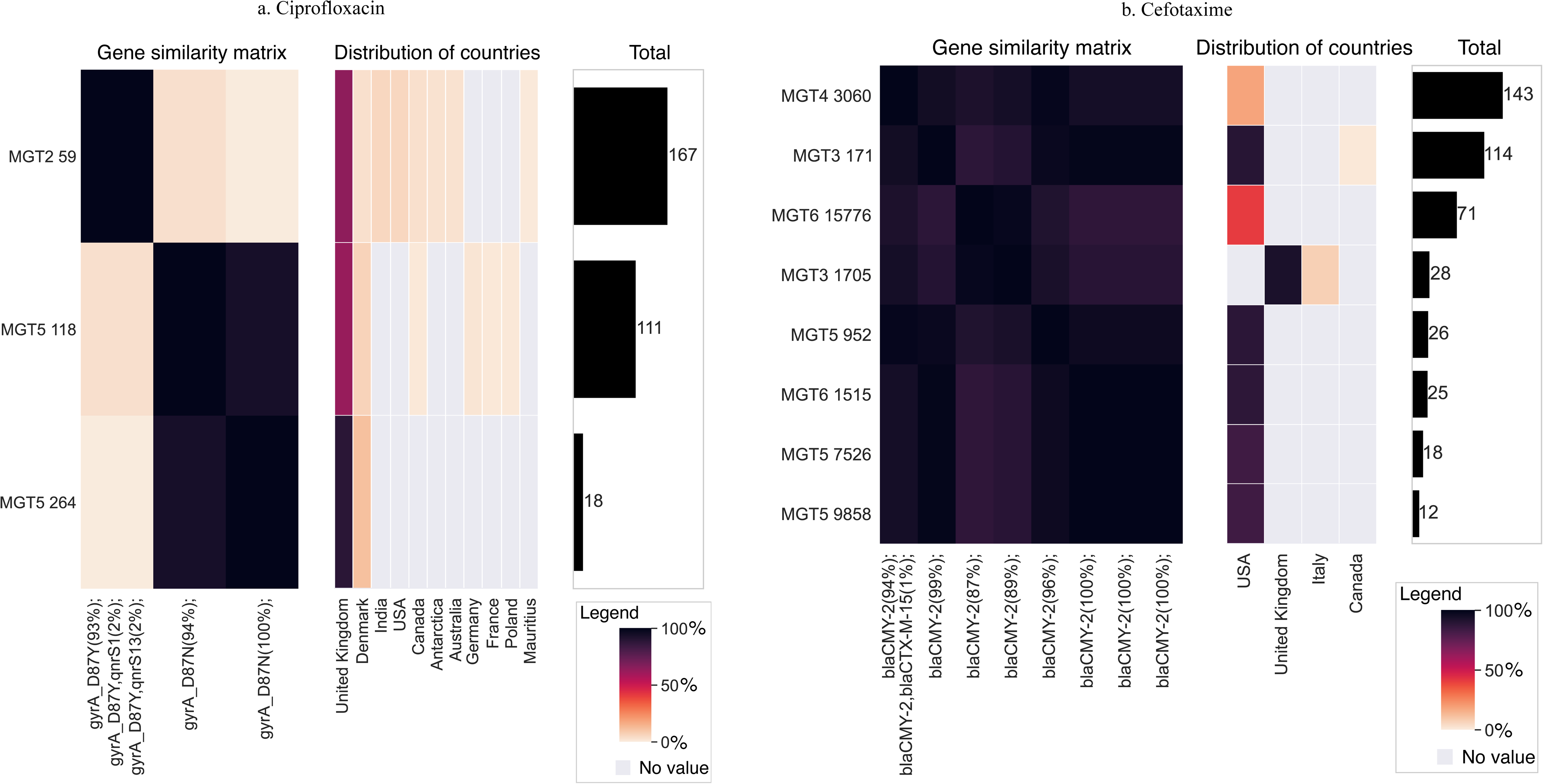
Non-susceptibility mechanism similarity matrix, country-distribution, and the total number of isolates, in non-susceptible MGT STs with at least 10 isolates in 2021-2022. a. Shows the three ciprofloxacin non-susceptible MGT STs, and b. shows eight cefotaxime non-susceptible MGT STs. The data shown here represents all isolates within the MGT STs, even those that are missing ‘year’ metadata in the country-distribution plots, or ‘country’ metadata in the non-susceptibility mechanism similarity matrix plots.

The diversity of ciprofloxacin non-susceptible isolates that were not classed within ciprofloxacin non-susceptible MGT STs is described in **Supplementary Results**.

### Cefotaxime

We identified 60 cefotaxime resistant MGT STs, which comprised 1,578 of 3,446 (46%) of cefotaxime resistant isolates (**Figure S8**), and included 96 susceptible isolates. In these 60 MGT STs, 923 isolates (55%) contained a year annotation, using which 22 MGT STs were identified as major STs in 2015 to 2022. The largest ST (MGT3 ST171) comprised 60 isolates within this timeframe, and the remaining averaged around 21 isolates per ST.

We temporally-clustered these 22 MGT STs to identify the temporal patterns within this set. We identified eight distinct temporal patterns (or clusters) within the 20 MGT STs (**Figure 3b**). Seven clusters had isolates from a single geographical majority, as follows: five clusters comprised isolates from USA (Clusters 2, 4, 5, 7 and 8), one cluster comprised isolates from UK (Cluster 1), and one cluster comprised isolates from Other (Cluster 6, with MGT3 ST136 containing isolates from Mexico, and MGT5 ST5868 containing isolates from China). One cluster comprised isolates from multiple countries (Cluster 3).

In 2021-2022 - the last two years with annotations available - eight MGT STs comprised atleast 10 isolates, namely MGT3 ST1705, MGT3 ST171, MGT4 ST3060, MGT5 ST9858, MGT5 ST7526, MGT5 ST952, MGT6 ST15776, and MGT6 ST1515. Seven of these eight MGT STs comprised isolates from USA, with one ST, namely MGT3 ST1705, comprising isolates mainly from UK (and Italy). All seven MGT STs comprising isolates from USA were within MGT1 ST19 (**Figure 4b**). MGT3 ST171, comprising isolates from UK, was within MGT1 ST34 with 89% of the isolates containing the ASSuT genotype (**Figure 4b**).

Only a partial set of isolates had source metadata available: of these the UK ST, MGT3 ST1705, contained a small number of isolates from pig. The USA MGT STs contained a small number of isolates from human, as well as either poultry sources (MGT3 ST171, MGT5 ST7526, MGT5 ST952, MGT6 ST1515), or cattle sources (MGT6 ST15776, MGT4 ST3060) and one did not have any source metadata (MGT5 ST9858). However, for these MGT STs, in the context of their lower level MGT STs (**Figure 4b**), a predominant non-human source was identifiable – with MGT3 ST171 and MGT3 ST8 comprising isolates from avian sources, and MGT3 ST23 comprising cattle and pig isolates, with further differentiation into a single non-human source at the MGT4 level, with MGT4 ST3060 and MGT4 ST31 only comprising cattle isolates.

With regards to the underlying mechanism of resistance, in these 60 MGT STs, there were eight gene or gene combinations conferring cefotaxime resistance (**Figure S10**). The gene *bla*_CMY-2_ was the most common mechanism for resistance, found within resistant MGT ST isolates found in 49 MGT STs. The eight MGT STs with atleast 10 isolates in 2021-2022, also contained this gene (**Figure 5b**). Other 11 MGT STs contained genes such as *bla*_CTX-M-9_, *bla*_CTX-M-15_ and *bla*_SHV-2A_.

The eight MGT STs that were found in 2021-2022 were associated with multiple plasmid replicon types (**Figure 4**). The UK MGT3 ST1705 was found associated with the IncI1-I(alpha) plasmids only, and the remaining seven USA MGT STs were all found associated with the IncC plasmids. Additional plasmids were found associated with various MGT STs, such as the IncX1 plasmid with MGT3 ST171 and MGT5 ST952, the Col(pHAD28) plasmid with MGT6 ST15776, MGT5 ST7526 and MGT6 ST1515, and the ColpVC plasmid with MGT5 ST9858 and MGT5 ST952.

The diversity of cefotaxime non-susceptible isolates that were not classed within cefotaxime non-susceptible MGT STs is described in **Supplementary Results**.

## Discussion

In this study, we analysed the predicted non-susceptibility to 15 antibiotic classes (including clinically relevant classes) in 65,000 publicly available STm genomes. The public dataset comprised data from multiple countries, and especially included high-quality surveillance genome data from the UK from 2015 onwards and the USA from 2019 onwards. We analysed the level of non-susceptibility by MGT STs, to identify trackable non-susceptible MGT STs, wherein at least ≥80% isolates within an ST were non-susceptible to a drug.

In the global dataset, nearly half of the isolates were predicted to be non-susceptible to one or more antibiotics, with the non-susceptible isolates being non-susceptible to between 1 to 12 antibiotics and averaging four antibiotics. In general, however, the presence of predicted non-susceptibility varied by location, time and in MGT1 STs. We observed that over 2015 to 2022, UK (with 60% non-susceptible isolates) had significantly higher non-susceptibility than USA (with 42% non-susceptible isolates). Furthermore, over the two timeframes considered (2015 to 2018, and 2019 to 2022), the proportion of non-susceptible isolates in UK significantly decreased in the second time-frame compared to the first, and vice-versa in USA. In both UK and USA, despite these changes in proportions, the average resistance score within the set of non-susceptible isolates was not significantly different. Thus, these changes were driven by changes in numbers of non-susceptible isolates, respectively, rather than by the overall acquisition or loss of resistance to additional classes. However, observing non-susceptibility at the level of individual antibiotics by locations, timeframes, or MGT1 STs, depicted these isolate sets both loosing as well as gaining resistance as shown in **Figures 1**, **S3** and **S4**. Furthermore, even the previously observed chromosomal patterns of genotypic resistance to the traditional phenotypic classes, such as ASSuT in ST34, and especially ACSSuT in DT104, were not very well preserved. As was seen in the results, in both DT104, and ST34, when resistance to additional classes was present, a fewer percentage of isolates comprised the previously observed genotypic-combination for the phenotypic-patterns, indicating altering genes for the phenotypic resistance. These results indicate the highly dynamic nature of AMR within STm.

Multiple factors affect the observed changes in resistance. Some examples include, proximity to other AMR species or sub-species [23], changing selection pressures, such as due to altered AMR prescriptions for different STm populations [24], and co-selection via other selective pressures [25]. Furthermore, sampling of the underlying data can also affect the observed changes in non-susceptibility. For example, surveillance was part of the UK’s national framework throughout 2015 to 2022 – which aimed to sequence all presumptive cases of salmonellosis. Hence the observed decrease in non-susceptibility over the two timeframes in UK likely represents a real reduction. Furthermore, this significant decrease in non-susceptibility could be attributed to various implemented antimicrobial stewardship programs in the UK [26, 27]. Although stewardship recommendations are also available in the USA [28, 29], the apparent significant increase in non-susceptibility in isolates from the USA can likely be partly attributed to implementation of a national *Salmonella* surveillance program with potentially more unbiased sampling. The dynamic AMR landscape highlights the importance of implementing continuous and consistent monitoring strategies.

Once, monitoring is set-up, the next steps involve the tracking of standardized strain and their AMR content [30] – and this is achievable via MGT. The MGT typing system, via assignment of nine STs for every isolate, provides a scalable, standardized, and multi-resolution typing approach for STm. As demonstrated in this paper, we utilized MGT to separate the dataset into smaller STs at higher resolution MGT-levels, and identified higher-resolution STs comprising mutually exclusive sets of isolates with very high non-susceptibility (minimally 80%) to an antibiotic. We identified 407 non-susceptible MGT STs for 14 of 15 antibiotics, spanning all MGT levels. Thus, through the increasing resolution of MGT levels, the appropriate resolution to describe a particular group of closely related isolates carrying a particular non-susceptibility determinant can be easily identified by sub-setting from the lowest-resolution to the highest-resolution levels. Furthermore, via this approach, we present a way to correlate elements of the core-genome, via sequence typing (i.e. MGT STs), to the accessory genome [31, 32] (e.g. AMR determinants that are horizontally acquired) - such that the clones or strains carrying non-susceptibility, can be easily identified, shared, and monitored. The utility of such a correlation is that an additional layer of confidence is added to the predicted non-susceptibility. For example, within a non-susceptible MGT ST, a 20% buffer of sensitive strains is allowed, which enables non-susceptibility to be inferred, even in the cases of erroneous or real sporadic loss of non-susceptibility (such as in the cases of low sequencing quality, assembly errors, or plasmid instability). This was exemplified in the potential underreporting of cefotaxime non-susceptibility in 14 of 437 isolates, representing an error rate of at most 3%, well within the 20% buffer. Thus, the inclusion of 20% of closely related isolates, i.e. isolates with the same ST, allow for such discrepancies to be tolerated.

MGT provides a naturally hierarchical setting to study related isolates, as exemplified in **Figure 4**. Isolates within any higher resolution MGT ST can be studied at lower resolution levels, where in isolates are grouped more broadly, and thus, the lower-resolution levels will often comprise a larger set of isolates. Often by additionally studying high-resolution MGT STs in STs in the context of lower resolution level STs that contain them, inferences about the epidemiology can be made, especially in the cases where isolate metadata are missing. These inferences can potentially be made regarding how isolates track over time, over location or in hosts – and such examples are described for both ciprofloxacin (see **Supplementary Discussion**) and cefotaxime non-susceptible MGT STs (below).

In the last two years of the dataset, 2021 and 2022, eight MGT STs were cefotaxime non-susceptible MGT STs (namely, MGT3 ST1705, MGT3 ST171, MGT4 ST3060, MGT6 ST15776, MGT5 ST9858, MGT5 ST7562, MGT5 ST952, MGT6 ST1515; see **Section Results Cefotaxime**). Epidemiologically, these eight MGT STs can be grouped into three distinct sets. MGT3 ST1705 was the only ST with isolates predominantly from UK and the only ST associated with pigs. This ST also had a distinctly associated plasmid replicon type, the IncI1-I(alpha). The other seven STs were from the USA. These STs were associated with either poultry or cattle, and had commonly the IncC plasmid replicon type. Both these plasmid replicon types have been previously associated with the carriage of the *bla*_CMY-2_ gene, and hence cefotaxime resistance [33, 34]. The unique association of distinct plasmids in in distinct MGT STs suggests that in both these groups acquisition of resistance to cefotaxime was independent. The seven MGT STs with isolates from USA were either from poultry or cattle. Interestingly, the poultry-associated non-susceptible MGT STs had high proportion of isolates with resistance to tetracyclines and sulfathiazoles, and the cattle associated non-susceptible MGT STs had a high proportion of isolates with resistance to tetracyclines, streptomycins, sulfathiazoles and chloramphenicol, further distinguishing the STs from the two sources based on their AMR profile. Thus, via our approach, identification of non-susceptible MGT STs for the same antibiotic, but associated with different reservoirs, can be easily identified and further tracked. This work here, clearly demonstrated how genomics-based precise tracking can be achieved for highly resistant strains.

Lastly, a brief discussion on the application of clustering approaches for the epidemiological surveillance of non-susceptible isolates via non-susceptible MGT STs is presented in the **Supplementary Discussion**.

## Conclusions

In this study, we paint a comprehensive picture of AMR in the publicly available global STm genome sequence data. In particular, we demonstrated the utility of integrating MGT to identify and enable precise tracking of non-susceptible MGT STs and temporal and spatial clustering to identify trends of non-susceptible MGT STs. The approaches presented here can be effectively applied to the standardised global comprehensive surveillance of AMR within STm, providing opportunities for early detection, data-driven decision making, targeted interventions, and measurements of effectiveness of interventions, and should be applicable to other organisms.

## Methods

### Genome assembly and assignment of MGT STs

We previously developed and maintain a database (MGTdb) for typing STm isolates using MGT [11, 21]. Briefly, the following pipeline forms part of MGTdb. Daily, any new isolates available in NCBI SRA [35], particularly Illumina paired-end sequenced, are imported into MGTdb. These, along with any isolates uploaded directly to MGTdb by users, are assembled using a pipeline which uses the SKESA algorithm [36]. From the assemblies, alleles are extracted which are utilized to type the isolates at all MGT levels [11]. The isolates imported from NCBI SRA are made publicly available, and the privacy of isolates uploaded by users can be set by the user to be either ‘public’ or ‘private’. Our dataset was composed of all public isolates within MGTdb, until August 31, 2023, that were typed at at-least one MGT level.

### AMR annotation

We utilized AbritAMR [37] (version 1.0.14) to assign the AMR profiles for each isolate. In particular, we ran AbritAMR, using the default options, on each assembly to identify the AMR determinants that were grouped into 14 classes by the tool. Following the identification of AMR determinants, we ran the ‘report’ option in AbritAMR, to obtain a predicted phenotypic interpretation for the 14 classes based on the mechanism of resistance. Of these 14 classes, two classes, cefotaxime (AmpC) and cefotaxime (ESBL), were combined into the one “cefotaxime” class, as both AmpC and ESBL were the mechanisms for resistance to the same antibiotic. Additionally, AbritAMR reported mechanisms for two additional classes colistin, and aminoglycoside resistance medicated via 16S rRNA methyltransferases (abbrev. aminoglycosides (Arm/Rmt)), however did not provide an interpretation. For simplicity, we interpreted the presence of any mechanism as ‘resistant’ for these two classes.

### Validation of AMR prediction

To validate the phenotypic prediction of resistance by AbritAMR, we compared isolates from NCBI Pathogen Detection [38] until 14 December, 2022, for which both phenotypic and Illumina-paired-end sequencing data were available. For either AST or genomics-based predictions, we considered two classes of resistance as either susceptible or non-susceptible for simplicity of comparison.

### Reliability of cefotaxime resistance prediction based on presence of *bla*_CMY-2_

To test reliability of *bla*_CMY-2_ annotations, isolates assigned eight MGT STs were reassembled using SPAdes [39] (version 3.15.5) with the option ‘--isolate’. Then, similarly as for SKESA, AbritAMR was run on the assemblies, followed by the ‘report’ option to generate an interpretation of resistance.

For comparison with an assembly free approach, we utilized KMA [40], a *k*-mer based matching algorithm given a template. For the template, from NCBI Pathogen Detection’s Reference Gene Catalog [41], we downloaded the *bla*_CMY-2_ nucleotide sequence (RefSeq ID: NG_048814.1). We indexed this sequence using KMA [40] (version 1.4.10) using the default settings. We then used KMA with default setting to match the reads to the template. The intact *bla*_CMY-2_ gene was inferred when coverage and identity of the reads to the matched template was 100% for both, otherwise the presence of a non-intact gene was inferred.

### Plasmid annotation

PlasmidFinder [42] (downloaded on 2023, June 4) using it’s default database was run on each isolate to infer the presence of plasmids via the presence of plasmid replicon sequences. The default threshold of coverage and identity of 60% and 95%, respectively, were utilized to identify the presence of plasmid replicon sequences.

### Non-susceptibility score (NS_s_)

For each isolate we calculated a non-susceptibility score (NS_s_). We represented an isolate’s susceptibility to an antibiotic as 0, and non-susceptibility as 1. A non-susceptibility score was then defined as the sum of the Boolean values across the 15 antibiotics. Given this score for each isolate, for any data subset (e.g. UK isolates between 2015 to 2018, or all MGT1 ST34 isolates), an average NS_s_, or average NS_s_ of non-susceptible isolates only, can be easily calculated as the average across those data subsets. Bootstraps using the NS_s_, for the datasets indicated in the results, was performed 9999 times in python (version 3.9.6) using the scipy.stats package (version 1.10.1).

The distribution of NS_s_ between different data subsets were compared using the Mann-Whitney *U* test [43] (with continuity set to False, and significance tested at a Bonferonni corrected *p*-value [44] as indicated in the results). The statistical test was performed using the scipy.stats package (version 1.10.1) in python3 (version 3.9.6).

### Comparison of non-susceptibility to individual antibiotics

To compare non-susceptibility between data subsets to individual antibiotics we used the Binomial test [45]. We considered susceptibility as 0, and resistance or intermediate resistance as 1, and compared the probability of observing either resistance or intermediate resistance, separately, in one data subset in comparison to another. Significance was tested at the Bonferroni corrected *p*-value as indicated in the results. The Binomial test implemented in the scipy.stats package (version 1.10.1) in python3 (version 3.9.6) was used.

### Identification of highly non-susceptible MGT STs

To identify non-susceptible MGT STs, i.e. MGT STs where at least 80% of isolates were non-susceptible to an antibiotic, the following procedure was implemented. For each antibiotic the MGT levels, from 1 to 9, were traversed, such that, at each level, the percentage of isolates with non-susceptibility to the antibiotic were calculated. Then isolates that were in STs with ≥80% non-susceptibility were removed from further analysis, and that MGT ST was marked as a non-susceptible MGT STs. This process was then repeated at the next level, with the remaining isolates. The isolates were removed to prevent an isolate from being counted within multiple non-susceptible MGT STs at different levels. This analysis was performed in python3 (version 3.9.6).

### Temporal clusters

For a given antibiotic, we standardized each non-susceptible MGT STs using the counts per years, as:

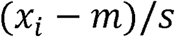

where *x_i_* is the number of isolates in an MGT ST in one year, *m* is the average count across all years, and *s* is the standard deviation. We used silhouette index [46] to initially identify the best number of clusters for a dataset, and then though manual inspection of the separation between the clusters, the final number of clusters were selected. Analyses were performed in R (version 4.3.1), using the bios2mds package [47] (version 1.2.3) to calculate the silhouette index, and the Mfuzz package [48] (version 2.60.0) to perform temporal clustering. We used a custom function in R to plot the resulting clusters. The clusters were manually rearranged based on visual inspection, such that the resulting clusters were ordered by when the initial weighted-average of the cluster was the highest.

### Non-susceptibility mechanism clustering

Within an MGT STs, for a given antibiotic, we calculated the percentage of isolates that were susceptible, and percentage of isolates with each non-susceptibility mechanism or combination of mechanisms. This set, comprising all non-susceptible mechanisms identified for a given antibiotic, and the empty set (indicating susceptibility to the antibiotic) was termed *M*. Then, we calculated a distance between any two MGT STs *i* and *j* as:

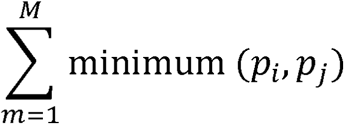

where, *p1* and *p2* are percentage of isolates in the two MGT STs, respectively, with *m* ∈ M.

Once this similarity matrix was calculated, we utilized the clustermap function in the seaborn library (version 0.12.2) in python3 (version 3.9.6), to perform average linkage clustering.

### MGT hierarchy

For each non-susceptible MGT ST, we displayed the MGT hierarchy. For each MGT ST, the lower level MGT ST was displayed if at least 50% of the isolates were assigned the same ST at the lower MGT level, otherwise the MGT ST was indicated with an ‘X’. Similarly, for each non-susceptible MGT ST, a plasmid replication type was annotated, if the annotation was present in at least 50% of the isolates within the ST. We also displayed maximally, the top five sources of the isolates within an ST, where available. However, sources were manually simplified to represent the closest associated animal. For example, “Human, stool” was simplified to “human”, “ground turkey” was simplified to “turkey. When the source was available for only a few isolates (<10) this was indicated by an asterisk.

The MGT hierarchy was generated in python3 (version 3.9.6) in dot format, which was then converted to a scalable vector graphic format using graphviz’s (version 8.0.5) hierarchical layout engine [49].

### World map

We simplified the “country” annotations of the isolates, e.g. “United States of America”, “United States, California” and “United States”, were all harmonized to “USA”. We then used the python3 (version 3.9.6) package, geopy (version 2.3.0), to identify the latitude and longitude for each country. The data were plotted on a world map using the packages ggplot2 [50] (version 3.4.4) and scatterpie (0.2.1) in R (version 4.3.1).

## Tables

See files Table 1.pdf, Table 2.pdf and Table 3.pdf.

