## Supplementary for "A dissection of the genomic antimicrobial resistance epidemiology of *Salmonella* Typhimurium"

### Supplementary Tables

Supplementary tables are provided in the file Supplementary Table 1.pdf, Supplementary Table 2.pdf and Supplementary Table 3.pdf.

### Supplementary Figure Legends

**Figure S1**. Sampling of data. a. Complete dataset. b-d. Isolates from UK, USA and Other, between 2015-2022.

**Figure S2**. Average non-susceptibility score in each year between 2015 to 2022 in UK, USA and Other. Additionally, for each graph, the bootstrap (9999 repeats), standard deviation, median of all data, and median of the grouped data as a 50^th^ percentile, is shown. The top row describes these statistics for all data in the three locations between 2015 to 2022, and the bottom row describes these statistics for non-susceptible isolates only.

**Figure S3**. Resistance to 15 antibiotics within the four most frequent MGT1 STs and the remaining STs. Blue outline indicates that the highlighted ST has the highest statistically significant antibiotic non-susceptibility between all MGT1 STs. Similarly, the yellow outline indicates significantly higher antibiotic non-susceptibility, however compared between ST19 and ST34 only, in the absence of an overall statistically significant indication between the two. The distributions of the presence or absence of non-susceptibility was assessed using the Binomial test, using a Bonferroni corrected *p*-value < 0.0001.

**Figure S4**. Non-susceptibility to 15 antibiotics in the UK and the USA, in 2015-2018 and 2019-2022, in three MGT1 STs: ST19, ST34 and the remaining STs. The drugs are arranged from left to right in descending order of non-susceptibility in the complete dataset. Significantly higher non-susceptibility in 2019-2022, compared to 2015-2018, in the UK and the USA, is indicated via a red upwards facing arrow, and significantly lower non-susceptibility is indicated via a green downwards facing arrow. Significantly higher non-susceptibility between the UK and the USA in 2019-2022 is indicated via a blue outline. Non-susceptibility to an antibiotic between two groups was tested using the Binomial test. Significance was assessed at the Bonferroni corrected *p*-value < 0.0002.

**Figure S5**. Yearly distribution of the phenotypic ASSuT pattern of resistance within ST34 isolates. a. Count of isolates per year, b. percentage of isolates per year.

**Figure S6**. Yearly distribution of phenotypic ACSSuT pattern of resistance within MGT3 ST11 (or DT104) isolates. a. Count of isolates per year, b. percentage of isolates per year.

**Figure S7**. Isolates included in non-susceptible MGT STs at different thresholds of minimal inclusion. The thresholds for minimally including non-susceptible isolates tested were 50%, 60%, 70%, 80% and 90%. At these percentage, a. shows percentage of non-susceptible isolates included in non-susceptible MGT STs. b. shows percentage of susceptible isolates included in non-susceptible MGTs.

**Figure S8**. Isolates included in non-susceptible MGT STs, where non-susceptible MGT STs are MGT STs where at least 80% isolates are non-susceptible to an antibiotic. In both sub-graphs, blue indicates non-susceptible isolates within such MGT STs, orange indicates non-susceptible isolates not within non-susceptible MGT STs, and green indicates susceptible isolates within non-susceptible MGT STs. a. Counts. b. Percentages, scaled to the total count of non-susceptible isolates for the given antibiotic.

**Figure S9**. Ciprofloxacin non-susceptibility mechanism similarity matrix, country-distribution and number of isolates in the complete set of ciprofloxacin non-susceptible MGT STs. The data shown here represents all isolates within the MGT STs, even those that are missing ‘year’ metadata in the country-distribution plots, or ‘country’ metadata in the non-susceptibility mechanism similarity matrix plots.

**Figure S10**. Cefotaxime non-susceptibility mechanism similarity matrix, country-distribution and number of isolates in the complete set of cefotaxime non-susceptible MGT STs. The data shown here represents all isolates within the MGT STs, even those that are missing ‘year’ metadata in the country-distribution plots, or ‘country’ metadata in the non-susceptibility mechanism similarity matrix plots.

**Figure S11**. Country and resistance-mechanism of 45 azithromycin resistant isolates in 2021 and 2022.

**Figure S12**. Country and resistance-mechanism of 31 colistin non-susceptible isolates in 2021 and 2022.

Supplementary figures are provided in the file Supplementary Figures.pdf.

### **Supplementary Code**

**Supplementary Code** – Scripts to enable all the analysis presented in this paper will be available at <https://github.com/Lanlab/AMR_analysis>.

### Supplementary Files

Supplementary File 1.txt – File describing 407 non-susceptible MGT STs.

**Supplementary Results**

### Non-susceptibility to any antibiotic

Within each location, yearly variation in non-susceptibility was observed (**Figure S2**). Upon comparison of the average presence or absence of non-susceptibility to any drug in isolates from UK and USA between 2015-2018 and 2019-2022, in the UK, the non-susceptibility significantly decreased from 64% (avg. NS_s_=2.5) to 55% (avg. NS_s_=2.3), whereas in USA it significantly increased from 42% (avg. NS_s_=1.4) to 48% (avg. NS_s_=1.6). However, when we compared the average NS_s_ of non-susceptible isolates only, there was no significant difference between the two timeframes for both UK (avg. NS_s_ of non-susceptible isolates only in 2015 to 2018=4, and in 2019-2022=4.1) and USA isolates (avg. NS_s_ of non-susceptible isolates only in 2015 to 2018=3.4, and in 2019-2022=3.4). Lastly, between isolates from the UK and USA in 2019-2022, UK had both a significantly higher proportion of non-susceptible isolates, as well as significantly higher average NS_s_ in non-susceptible isolates only.

### Comparison of non-susceptibility to individual antibiotics between UK and USA

We also compared the resistance to individual antibiotics between the UK and the USA in 2019-2022 (**Figure 1**). The UK had significantly higher predicted non-susceptibility to eight antibiotics, namely streptomycin (higher by 23%), ampicillin (by 25%), chloramphenicol (by 8%), trimethoprim (by 8%), trimethoprim-sulphathiazole (by 7%), kanamycin (by 1.3%), azithromycin (by 0.5%), and meropenem (by 0.1%), and the USA had higher non-susceptibility to only two antibiotics, namely, cefotaxime (higher by 5%) and gentamicin (by 5%). Resistance to ciprofloxacin was not significantly different between UK and USA, however intermediate resistance was higher in UK (by 4%).

### Non-susceptibility within MGT1 (MLST) STs for individual antibiotics

Differences in non-susceptibility to individual antibiotics was also compared between STs **(Figure S3**). As expected, between ST34 and ST19 isolates, ST34 had higher non-susceptibility to most drugs (11 of 15), including the currently prescribed antibiotics, ciprofloxacin (higher resistance by 1%, higher intermediate resistance by 2%), azithromycin (by 2%), and the last-line antibiotic colistin (by 2%). ST19 isolates had higher non-susceptibility to three drugs, including the currently prescribed antibiotic cefotaxime (higher by 2%).

Non-susceptibility to each antibiotic in each ST were compared between the UK and the USA for 2019-2022 to understand if variability was present within an STs by location (**Figure S4**). ST19 isolates from the UK had significantly higher non-susceptibility to five drugs, including ampicillin (higher by 10%) and ciprofloxacin (higher intermediate resistance by 7%). ST19 in USA also had higher non-susceptibility to six drugs, including cefotaxime (higher by 6%). ST34 isolates from the UK had higher non-susceptibility to seven drugs, and the USA ST34 had higher non-susceptibility to five drugs including ciprofloxacin (higher resistance by 1% and higher intermediate resistance by 12%), cefotaxime (higher by 4%) and colistin (by 5%).

Distribution of the ASSuT resistance pattern

We determined the proportion of ST34 isolates which had the previously observed ASSuT phenotypic resistance pattern, and the corresponding genes (‘*bla*_TEM-1_, *aph(3'')-Ib* and *aph(6)-Id*, *sul2*, and *tet*(B)). In the set of 9,122 ST34 isolates, 70% percent had the phenotypic ASSuT resistance pattern, with 53% comprising resistance to exactly these four classes, and 17% with resistance to additional classes (i.e. ASSuT+) (**Figure S5**). Resistance to trimethoprim, and hence trimethoprim-sulphathiazole, was the second most frequent resistance, present in 58% of ASSuT+ isolates. Of the ST34 isolates with the ASSuT or the ASSuT+ resistance pattern, 90%, had the exact genotypic combination of ‘*bla*_TEM-1_, *aph(3'')-Ib* and *aph(6)-Id*), *sul2*, and *tet*(B)’. In particular, this exact genotypic combination was found in 99% of isolates with ASSuT resistance, and 62% of isolates with ASSuT+ resistance.

Distribution of the ACSSuT resistance pattern

Similarly, we determined the proportion of DT104 (or MGT3 ST11) isolates, which had the previously observed ACSSuT phenotypic resistance pattern, and the corresponding genes (*tet*(G*)*, *tet*R, *aadA2*, *bla*_CARB-2_, *sul1*, *floR*). The tetracycline resistance gene *tet*R is not present in the AMRFinderPlus database [1], and hence its identification has not been included in this analysis. In the set of 4,852 MGT3 ST11 isolates, only 15% had the predicted phenotypic resistance to the five classes, with 7% comprising resistance to exactly these five classes, and 8% with resistance to additional classes (i.e. ACSSuT+) (**Figure S6**). The absence of resistance to sulfathiazole was most common, with sulfathiazole susceptibility observed in 81% of isolates with resistance to atleast one other class from ACSSuT. Lastly, in the isolates with ACSSuT or ACSSuT+, 59% had the exact genotypic combination ‘*tet*(G*)*, *aadA2*, *bla*_CARB-2_, *sul1*, *floR*’. In particular, this exact genotypic combination was found in 90% of isolates with ACSSuT, and 31% of isolates with ACSSuT+ resistance.

Selection of the 80% cutoff to define non-susceptible MGT STs

The 80% cutoff was chosen to define non-susceptible MGT STs, i.e. a non-susceptible MGT ST must consist of at least 80% isolates that are non-susceptible to an antibiotic. This cutoff was chosen, because at this threshold the recall (i.e. the number of non-susceptible isolates included within non-susceptible MGT STs) was considerably higher compared to 90% for several antibiotics (particularly tetracycline, streptomycin, ampicillin and sulfathiazole), and at lower thresholds (70%, 60% and 50%) further large differences in recall were not seen (**Figure S7a**). However, precision (i.e. the number of susceptible isolates included within non-susceptible MGT STs) decreased at lower thresholds (**Figure S7b**).

### Highly non-susceptible MGT STs to a given antibiotic

**Azithromycin**

We identified three azithromycin resistant MGT STs, which comprised 78 of 629

(12%) of azithromycin resistant isolates (**Figure S8**), and included three susceptible isolates. Of these three MGT STs, only isolates in one ST, namely MGT5 ST4133, consisting of 15 isolate, had year metadata. These MGT5 ST4133 isolates were annotated as isolated in 2017, from mostly the UK. Three isolates were annotated as originating from human. This ST was within MGT1 ST34, and all resistant isolates within this ST comprised the *mph(A)* gene. Of the other two STs, MGT5 ST5010 was within ST102 and all resistant isolates comprised the *acrB* R717Q mutation, and MGT5 ST3370 was within ST313 and comprised the *mph(A)* gene.

The majority of isolates predicted to be azithromycin resistant (88%, 551 of 629) were not within resistant MGT STs. These isolates were found in all locations: UK (181), USA (95) and Other (168), with the remaining missing country annotation. These resistant isolates were distributed across 12 MGT1 STs, 44 MGT2 STs and 131 MGT3 STs. These isolates ranged from 1988 to 2022, with 45 isolates isolated in 2021-2022. These 551 resistant isolates had 28 different resistance genes, partial genes, mutations, or combinations of these, with the *mph(A)* gene being the most frequent in 40% of isolates. In 2021-2022, isolates with *mph*(A) were found in multiple countries, whereas isolates with *mef*(B) were only isolated in the UK (**Figure S11**).

**Colistin**

We identified four colistin non-susceptible MGT STs, which comprised 101 of 423 (24%) of colistin non-susceptible isolates (**Figure S8**), and included 13 susceptible isolates. In these four MGT STs, 110 isolates (97%) contained a year annotation. None of these four STs were major STs in the last two years of the dataset (i.e. 2021-2022). In particular, within the 2015-2022 timeframe, MGT3 ST1583 comprised isolates between 2015 to 2017 from mostly Australia, MGT5 ST5868 comprised isolates isolated in 2020 from predominantly China, MGT5 ST645 comprised isolates between 2015 to 2019 with one isolate from 2022. This ST comprised isolates mainly from UK, with other countries annotated as Belgium and Canada. Lastly, MGT6 ST957 comprised isolates from 2015 and 2016 from the UK. All four STs were within MGT1 ST19. Three of these STs (namely MGT3 ST1583, MGT5 ST645 and MGT6 ST957) comprised isolates containing the *mcr-9* gene for resistance to colistin, and one (MGT5 ST5868) comprised isolates containing the *mcr-1.1* gene for resistance to colistin.

The majority of isolates predicted to be colistin non-susceptible (76%, 322 of 423) were not within non-susceptible MGT STs. These isolates were found in all locations: UK (97), USA (49) and Other (152), with the remaining missing country annotation. These non-susceptible isolates were distributed across 8 MGT1 STs (with 68% in ST34), 16 MGT2 STs and 69 MGT3 STs. These isolates ranged from 1984 to 2022, with 31 isolates isolated in 2021-2022. These non-susceptible isolates had 21 different resistance genes, partial genes, mutations, or combinations of these, with *mcr-1.1* and *mcr-9* being equally frequent at 34% each. In 2021-2022, most colistin non-susceptible isolates were isolated from USA (22 of 31) and comprised *mcr-9* as the most frequent resistance mechanism (in 21 isolates) (**Figure S12**).

**Carbapenems**

In the entire dataset, resistance to carbapenems was very low, with only 14 resistant isolates. Meropenem resistant MGT STs were not identified. Of the 14 isolates, 7 isolates were isolated in 2021-2022. The gene *bla*_OXA-48_ was the most frequent resistance mechanism, present in 12 isolates, and the only mechanism in 2021-2022. In 2021-2022, five meropenem resistant isolates were isolated from UK, two from South Africa, with the remaining missing country annotations. These isolates were distributed in three MGT1 STs.

**Ciprofloxacin non-susceptible isolates not within non-susceptible MGT STs**

The majority of isolates predicted to be ciprofloxacin resistant (59%, 2,636) were not within ciprofloxacin resistant MGT STs. These isolates were found in all locations: UK (797), USA (399) and Other (649) with the remaining missing country annotation. These resistant isolates were highly genomically diverse, they had 104 different ciprofloxacin non-susceptibility genes, partial genes, mutations, or combinations of these, and were distributed across 36 different MGT1 STs, and 453 MGT3 STs.

**Cefotaxime non-susceptible isolates not within non-susceptible MGT STs**

Half of the isolates predicted to be cefotaxime resistant (54%, 1,868) were not within resistant MGT STs. These isolates were found in all locations: USA (936), UK (158) and Other (311) with the remaining missing country annotation. These resistant isolates had 44 different resistance genes, partial genes, or combinations of these, with the gene *bla*_CMY-2_ being the most frequent in 74%. These resistant isolates were distributed across 24 MGT1 STs, with again MGT1 ST19 being the most common (in 72% of isolates).

Reliability of prediction

We validated the prediction pipeline, by comparing the non-susceptibility predicted by our methods with data from NCBI pathogen detection. We found an overall accuracy of 98.6% (**Table S3**).

For ciprofloxacin, mutations in the gene encoding DNA gyrase subunit A (or *gyrA*) were found associated with non-susceptibility in recent years (2021 to 2022). This gene is part of the core-genome of STm (MGT8), and additionally forms part of the MGT7 scheme [2]. As this gene is part of the core genome, mutations in this gene will result in new STs at both MGT7 and MGT8 levels. However, in this analysis, lower levels (an MGT2, and two MGT5 STs) were sufficient to identify the non-susceptible isolates.

For cefotaxime, the presence of beta lactamase CMY-2 (or *bla*_CMY-2_) was the most frequent mechanism for resistance in recent years. This gene is does not form part of any MGT scheme [3], and it’s carriage has been observed on various plasmids in *Salmonella* [4]. We tested if the assembly approach utilized, or the non-susceptibility prediction pipeline used, had any effect on the identification of the presence of this gene.

To test if the assembly algorithm had any effect on non-susceptibility prediction, we compared the prediction using isolates within the eight MGT STs with atleast 10 isolates from 2021 to 2022 (see **Section Results Cefotaxime** for details). These MGT STs consisted of 437 isolates wherein 20 isolates were predicted to be susceptible to cefotaxime. We compared the predictions obtained using AbritAMR on two different assemblies for each isolate that were generated using two different tools, namely, SKESA [5] and SPAdes [6]. SKESA also forms a part of the MGT pipeline [2]. All isolates identified as resistant using SKESA assemblies, were also identified as resistant using SPAdes assemblies – however when using SPAdes, an additional 10 isolates were predicted as resistant, with five comprising the intact *bla*_CMY-2_ gene and five comprising the non-intact gene (albeit missing less that 10% of the sequence).

We then utilized KMA[7], a *k*-mer based approach for matching and aligning raw-reads to a query sequence, to determine if an assembly-free approach performs better for non-susceptibility prediction. Using KMA, an additional four isolates were identified as comprising the *bla*_CMY-2_ gene, where in one isolate comprised the intact gene, and three comprised the non-intact gene. In total, five isolates of 14, comprised the non-intact gene. Finally, six isolates were consistently identified as susceptible using all three methods.

The AbritAMR tool for non-susceptibility prediction, at its core, relies on the identification of the presence of non-susceptible mechanisms via AMRFinderPlus [1], and adds an interpretation of non-susceptibility based on the predicted mechanisms. Using either SKESA or SPAdes assemblies, AMRFinderPlus was able to predict the presence of the *bla*_CMY-2_ gene in all 14 discrepant genomes, however, when between 60-90% of the sequence was missing, AbritAMR assigned an interpretation of ‘susceptible’. Despite this, a potential underreporting of atmost 14 genomes represents an error rate of atmost 3%, comparable with the overall accuracy.

### Supplementary discussion

MGT enabled the identification of geographically restricted non-susceptible STs

In the last two years of the dataset, 2021 and 2022, two MGT5 STs were ciprofloxacin non-susceptible MGT STs (MGT5 ST118 and MGT5 ST264, see **Section Results Ciprofloxacin**). Both these STs are closely related, both isolated from primarily human, and sparsely bovine, from mostly UK. Hierarchically, both these STs were within the same MGT4 ST, and within MGT3 ST11 or DT104 at the MGT3 level. Although DT104 is considered a past pandemic clone, replaced by ST34 [8], it is yet present, and causing disease in humans. Thus, MGT STs enables the identification of ciprofloxacin-resistant trackable types within DT104 that are geographically restricted. In UK, decreased susceptibility to ciprofloxacin due to *gyrA* D87N has been observed in DT104 isolates from dairy cattle since at least 2000 [9]. Furthermore, as certain *Salmonella* infections may persist for life within an animal [10], or may persist in herds via various farming practices [11], this suggests that the previously implicated dairy farms may be a potential source of currently observed infections.

Clustering approaches can be easily adapted for epidemiological surveillance of non-susceptible isolates

We clustered the non-susceptible MGT STs in two ways, firstly, temporally, using the counts of isolates in year, and secondly, based on the similarity of the non-susceptibility mechanisms. Temporal clustering enables the temporal-profile of an STs to be compared with other STs, and also enables the identification of dominant temporal trends in the dataset. Similarly, the clustering of genetic mechanisms of non-susceptibility also enables the dominant mechanisms to be identified in a dataset. For example, for cefotaxime-resistance, within the entire dataset, although a number of mechanisms with resistant to the same drug have arisen in the past, only one mechanism *bla*_CMY-2_, has been observed in the last two years of the dataset. Although the application of such unsupervised clustering approaches are common in other life-sciences domain (such as time series proteomic measurements [12]), to our knowledge its application to epidemiological data, by considering MGT STs as multivariate, is novel. These approaches however provided a natural setting to understand the trends of non-susceptibility within a dataset and enabled comparison between different MGT STs with different abundances. Thus, via this work we presented an approach for applying clustering methods for epidemiological surveillance of non-susceptible isolates*.*

Whereas the non-susceptibility mechanism-based clustering relies on the extraction of information from the genome-sequence, temporal clustering relies on the annotation of ‘time’. Thus, the interpretation of an observed temporal-trend depends on both annotation of time, as well as the underlying sampling, i.e. if the underlying sampling is unbiased and continuous, the observed trend of an ST is a true representation of its epidemiology. Again, this highlights the importance of sampling for epidemiological inferences.
