## Supplementary Figures for "A dissection of the genomic antimicrobial resistance epidemiology of *Salmonella* Typhimurium"

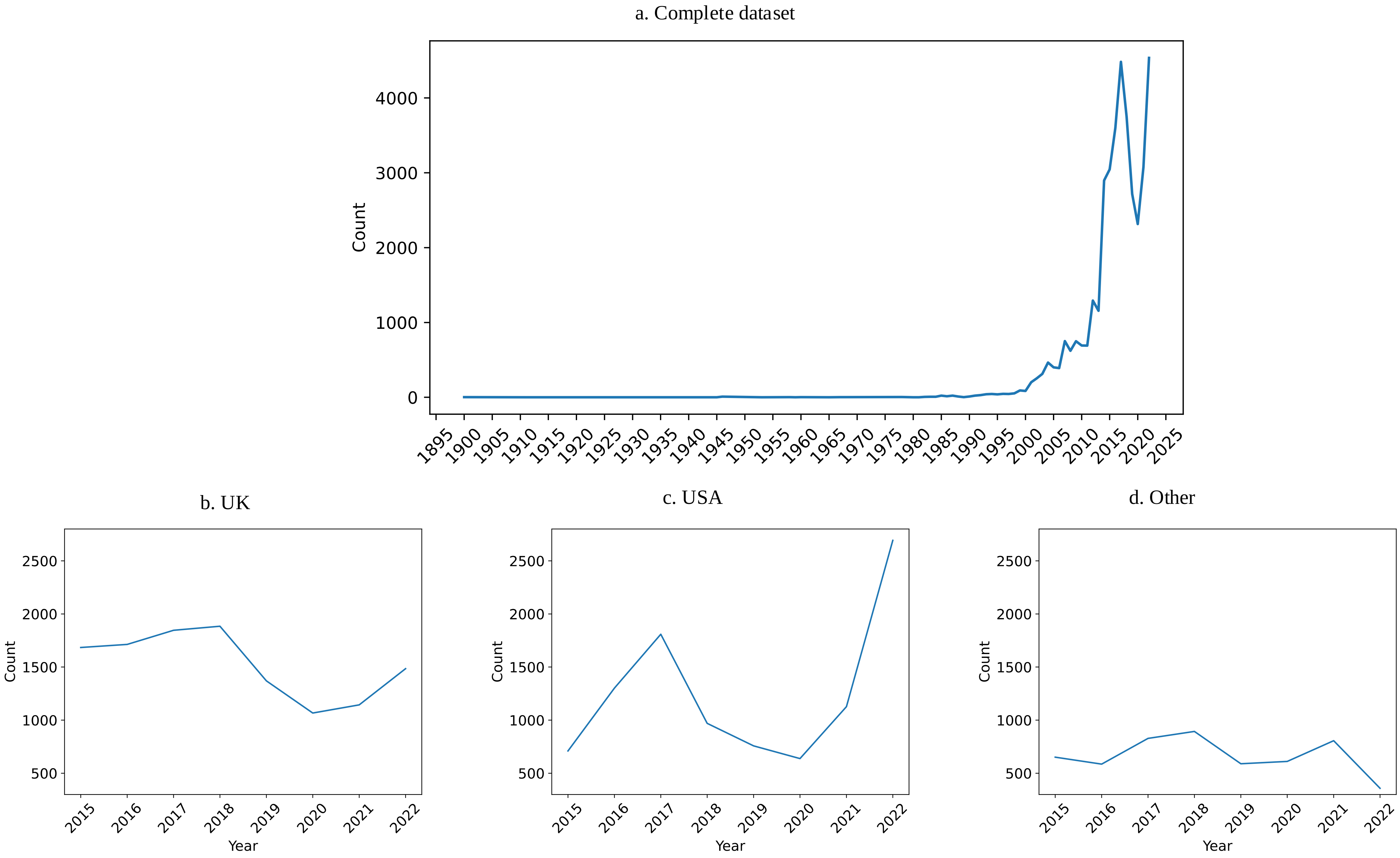


**Figure S1.**

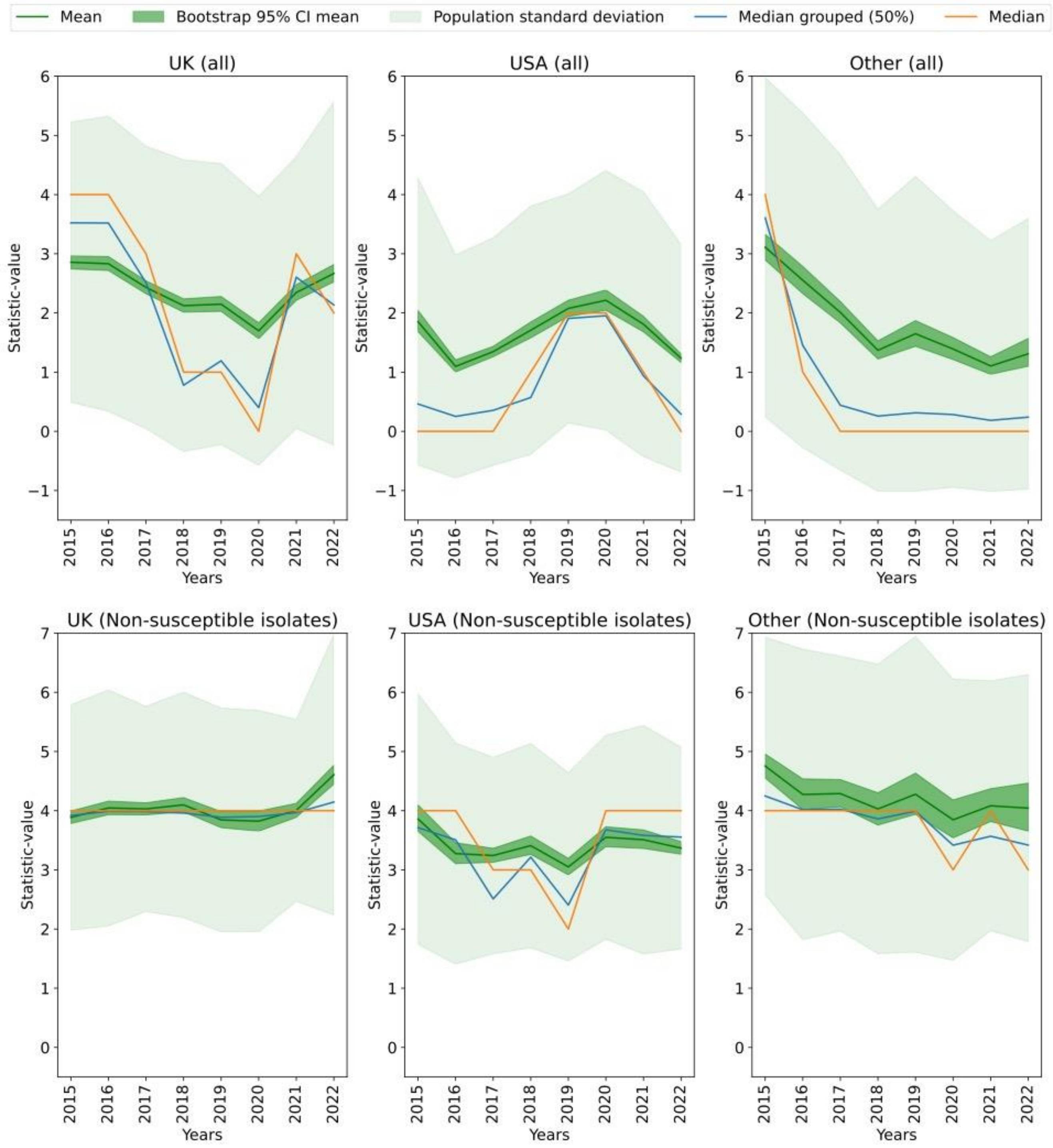

Mean
Bootstrap 95% CI mean
Population standard deviation
Median grouped (50%)
Median
UK (all)
USA (all)
Other (all)
UK (Non-susceptible isolates)
USA (Non-susceptible isolates)
Other (Non-susceptible isolates)
Statistic-value
Years


**Figure S2**.

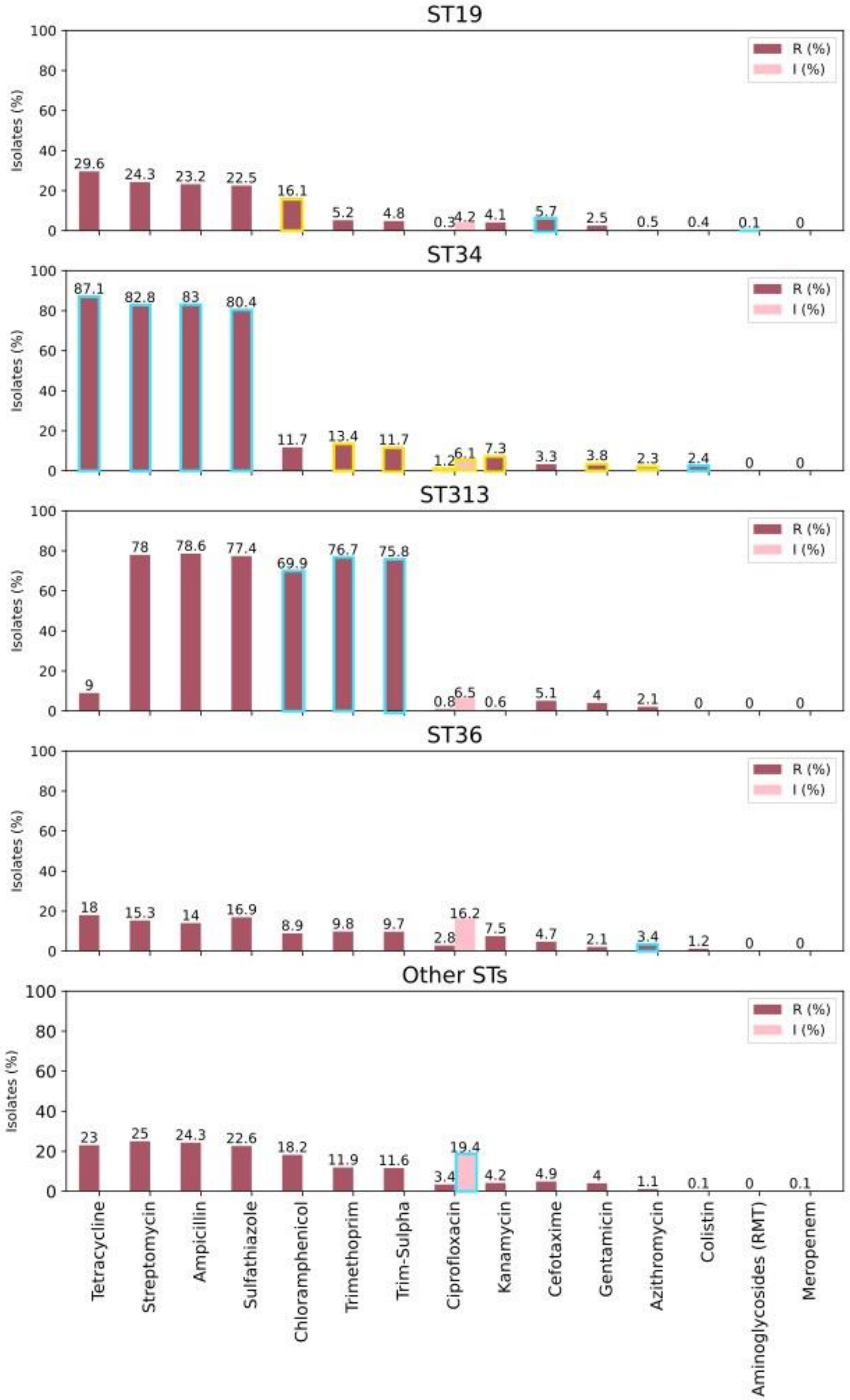
ST19
ST34
ST313
ST36
Other STs
Isolates (%)
R (%)
I (%)
Tetracycline
Streptomycin
Ampicillin
Sulfathiazole
Chloramphenicol
Trimethoprim
Trim-Sulpha
Ciprofloxacin
Kanamycin
Cefotaxime
Gentamicin
Azithromycin
Colistin
Aminoglycosides (RMT)
Meropenem

**Figure S3**.

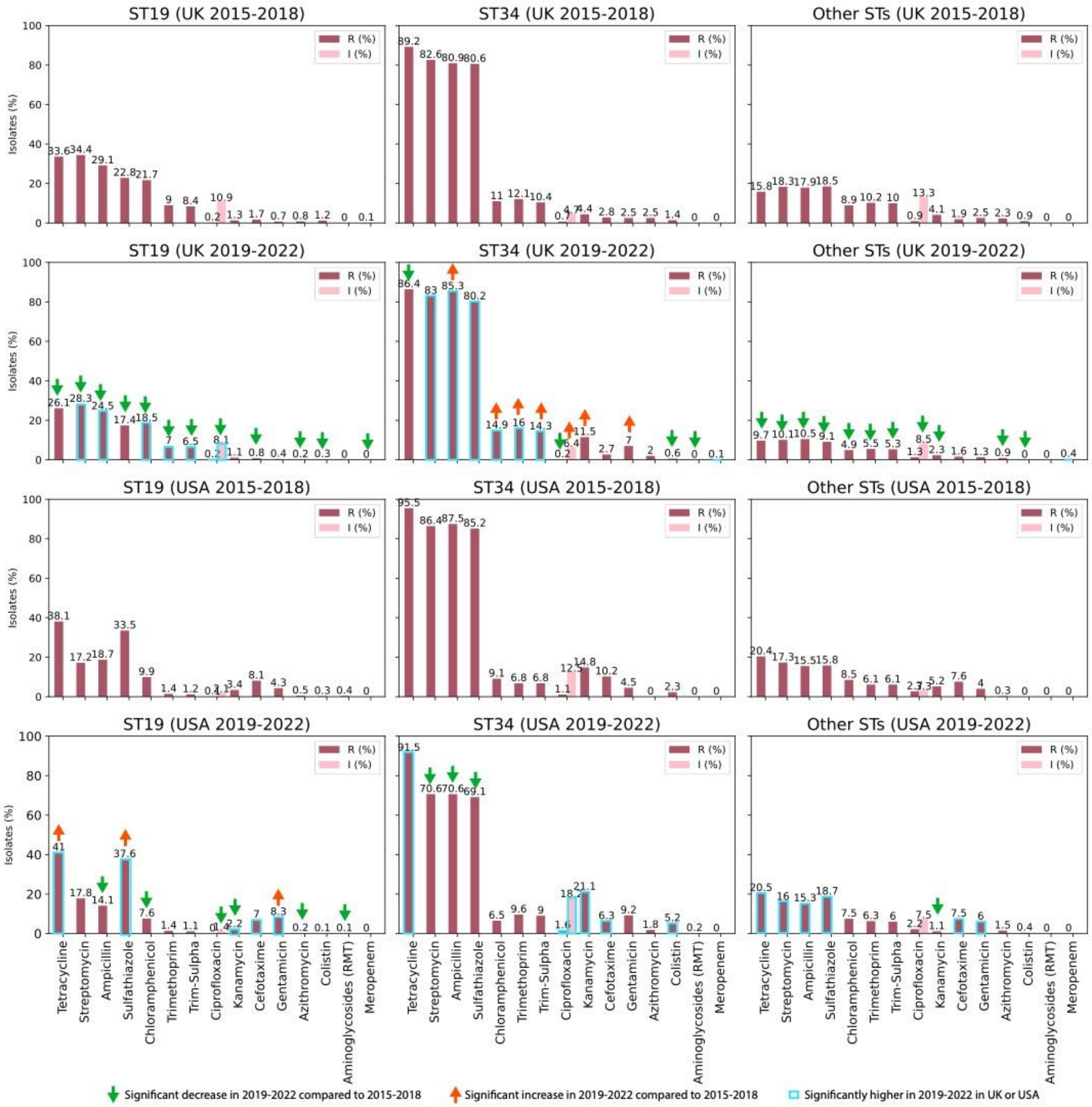
ST19 (UK 2015-2018)
ST34 (UK 2015-2018)
Other STs (UK 2015-2018)
ST19 (UK 2019-2022)
ST34 (UK 2019-2022)
Other STs (UK 2019-2022)
ST19 (USA 2015-2018)
ST34 (USA 2015-2018)
Other STs (USA 2015-2018)
ST19 (USA 2019-2022)
ST34 (USA 2019-2022)
Other STs (USA 2019-2022)
R (%)
I (%)
Isolates (%)
Tetracycline
Streptomycin
Ampicillin
Sulfathiazole
Chloramphenicol
Trimethoprim
Trim-Sulpha
Ciprofloxacin
Kanamycin
Cefotaxime
Gentamicin
Azithromycin
Colistin
Aminoglycosides (RMT)
Meropenem
Significant decrease in 2019-2022 compared to 2015-2018
Significant increase in 2019-2022 compared to 2015-2018
Significantly higher in 2019-2022 in UK or USA

**Figure S4.**

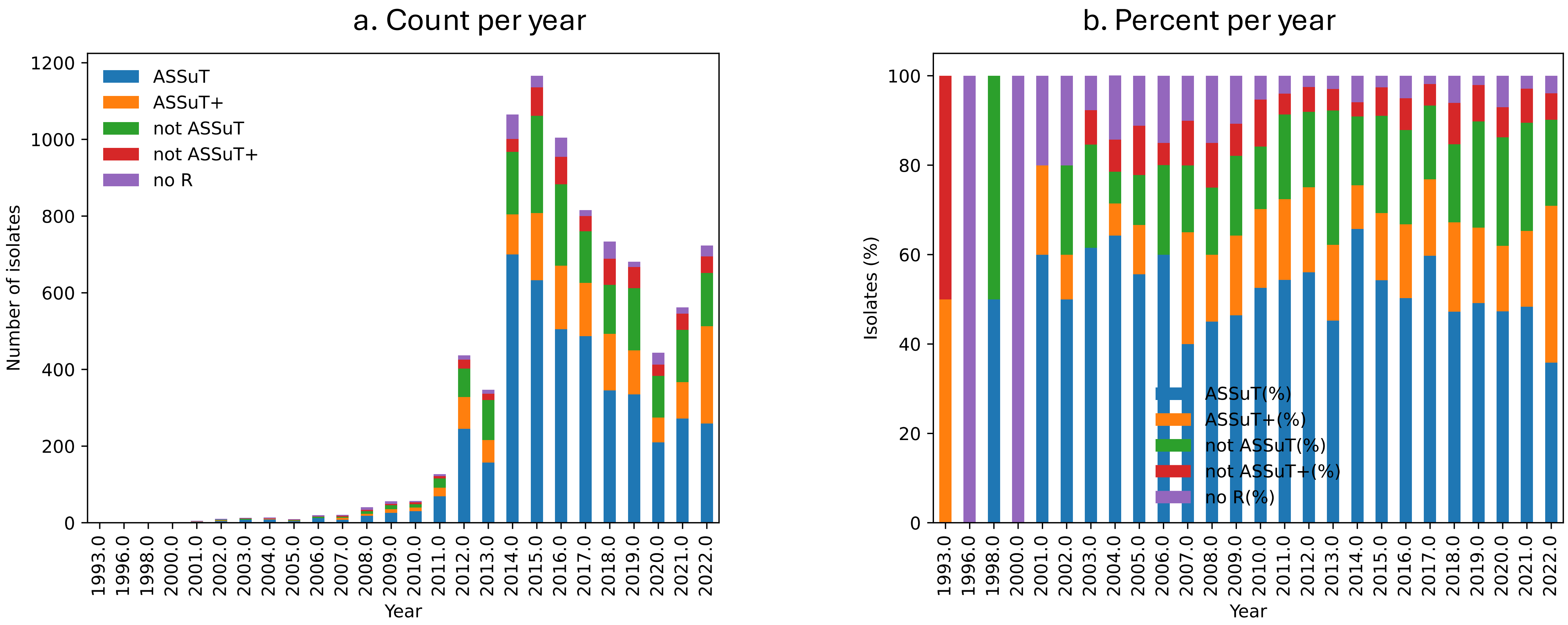

a. Count per year
ASSuT
ASSuT+
not ASSuT
not ASSuT+
no R
Number of isolates
0
200
400
600
800
1000
1200
Year
1993.0
1996.0
1998.0
2000.0
2001.0
2002.0
2003.0
2004.0
2005.0
2006.0
2007.0
2008.0
2009.0
2010.0
2011.0
2012.0
2013.0
2014.0
2015.0
2016.0
2017.0
2018.0
2019.0
2020.0
2021.0
2022.0
b. Percent per year
ASSuT(%)
ASSuT+(%)
not ASSuT(%)
not ASSuT+(%)
no R(%)
Isolates (%)
0
20
40
60
80
100
Year


**Figure S5**.

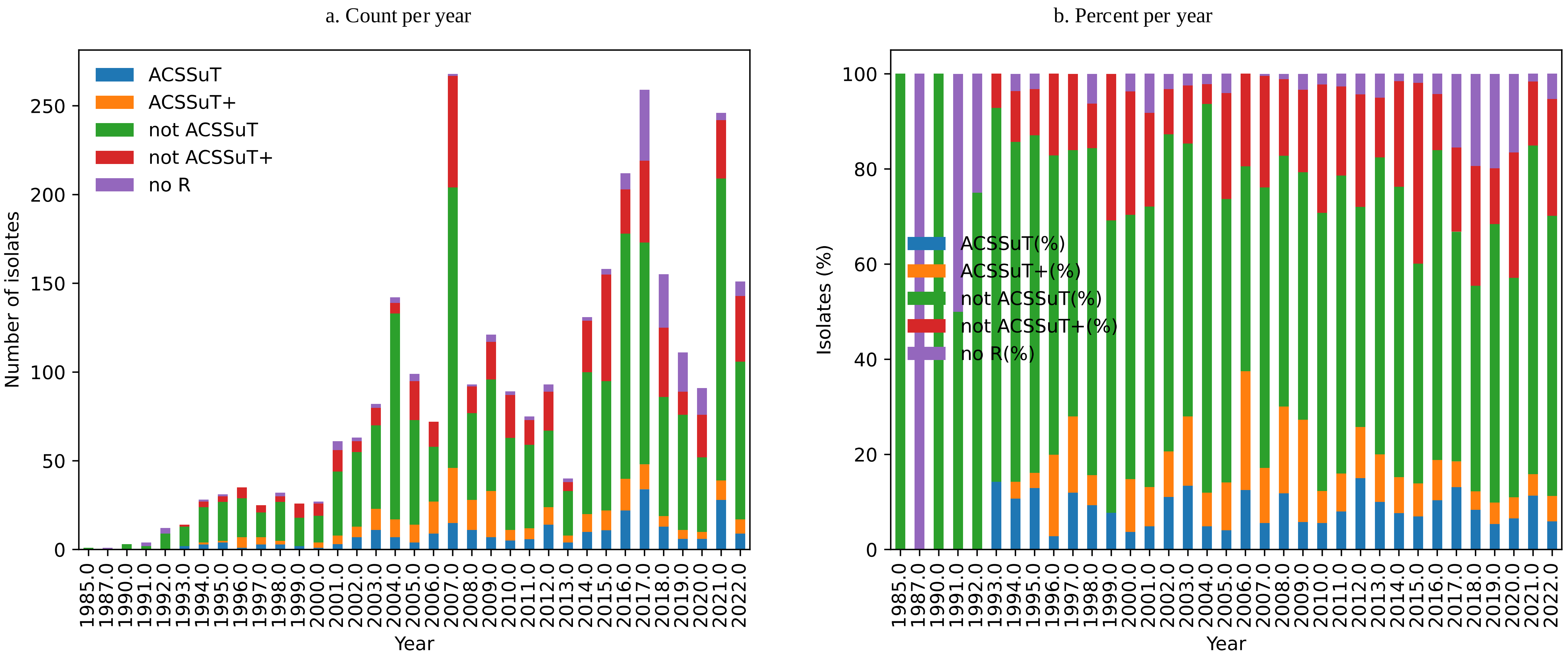

a. Count per year
ACSSuT
ACSSuT+
not ACSSuT
not ACSSuT+
no R
Number of isolates
Year
b. Percent per year
ACSSuT(%)
ACSSuT+(%)
not ACSSuT(%)
not ACSSuT+(%)
no R(%)
Isolates (%)
Year


**Figure S6**.

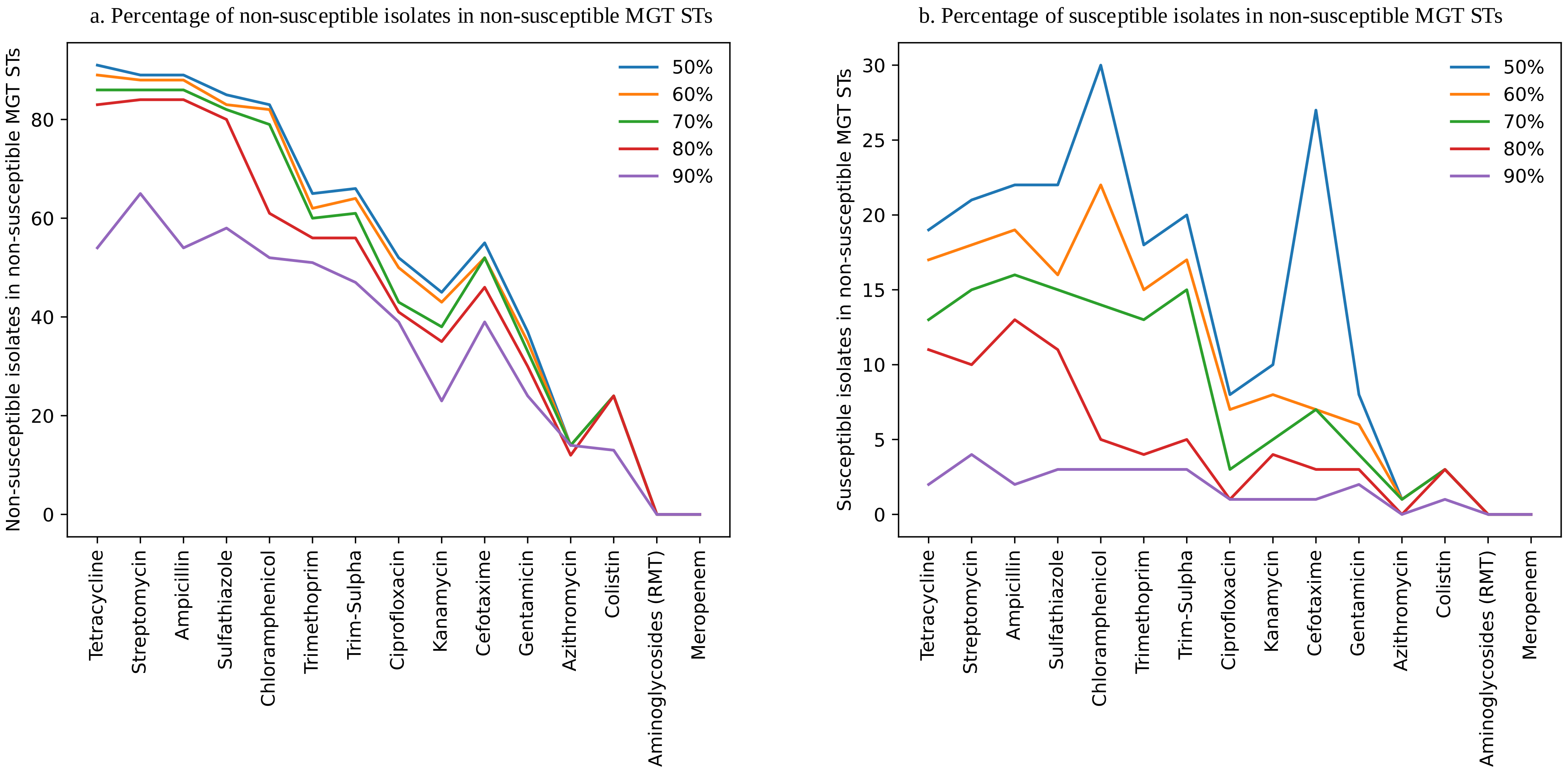
a. Percentage of non-susceptible isolates in non-susceptible MGT STs
Non-susceptible isolates in non-susceptible MGT STs
0
20
40
60
80
50%
60%
70%
80%
90%
Tetracycline
Streptomycin
Ampicillin
Sulfathiazole
Chloramphenicol
Trimethoprim
Trim-Sulpha
Ciprofloxacin
Kanamycin
Cefotaxime
Gentamicin
Azithromycin
Colistin
Aminoglycosides (RMT)
Meropenem
b. Percentage of susceptible isolates in non-susceptible MGT STs
Susceptible isolates in non-susceptible MGT STs
0
5
10
15
20
25
30
50%
60%
70%
80%
90%
Tetracycline
Streptomycin
Ampicillin
Sulfathiazole
Chloramphenicol
Trimethoprim
Trim-Sulpha
Ciprofloxacin
Kanamycin
Cefotaxime
Gentamicin
Azithromycin
Colistin
Aminoglycosides (RMT)
Meropenem

**Figure S7**.

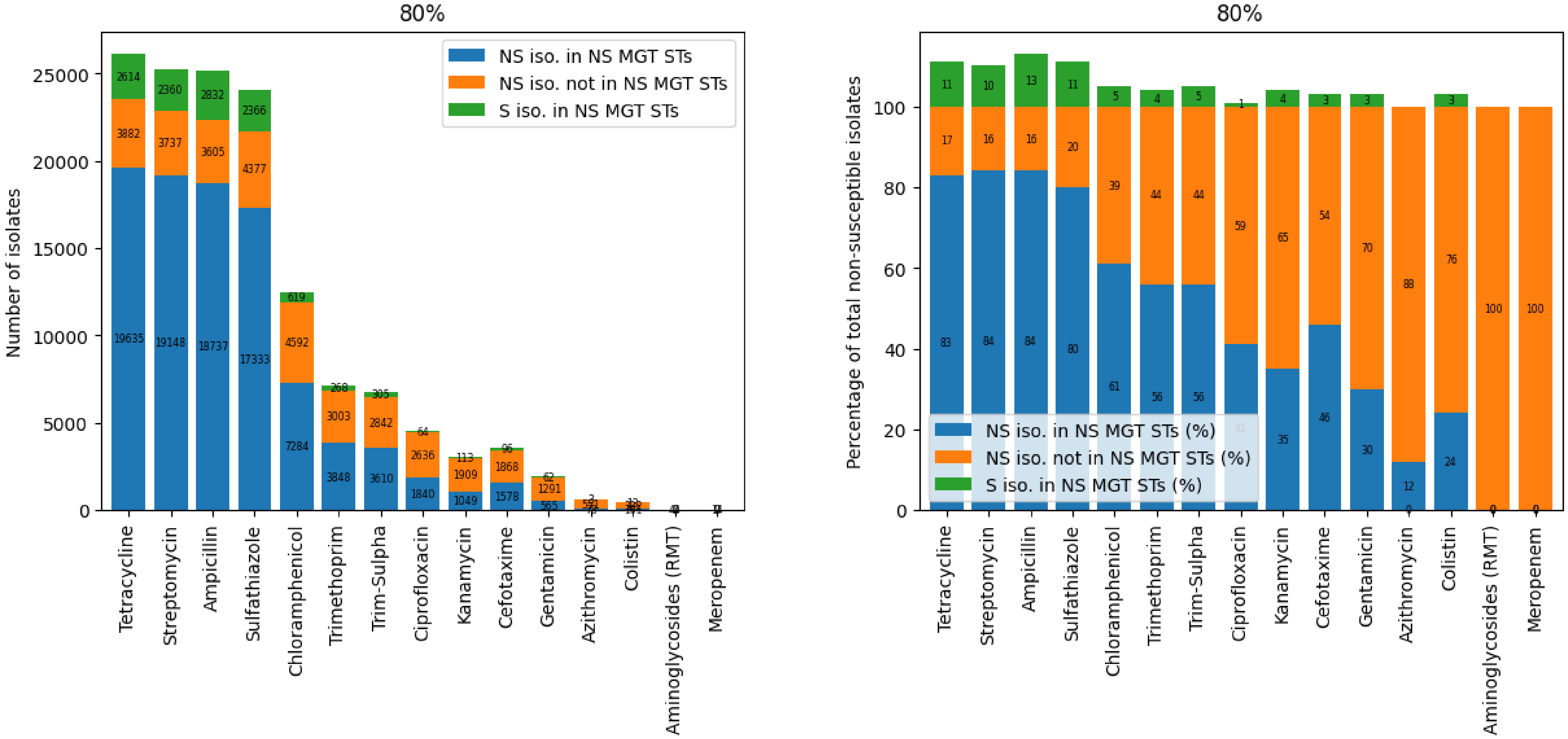


a. Count per year

b. Percent per year

**Figure S8**.

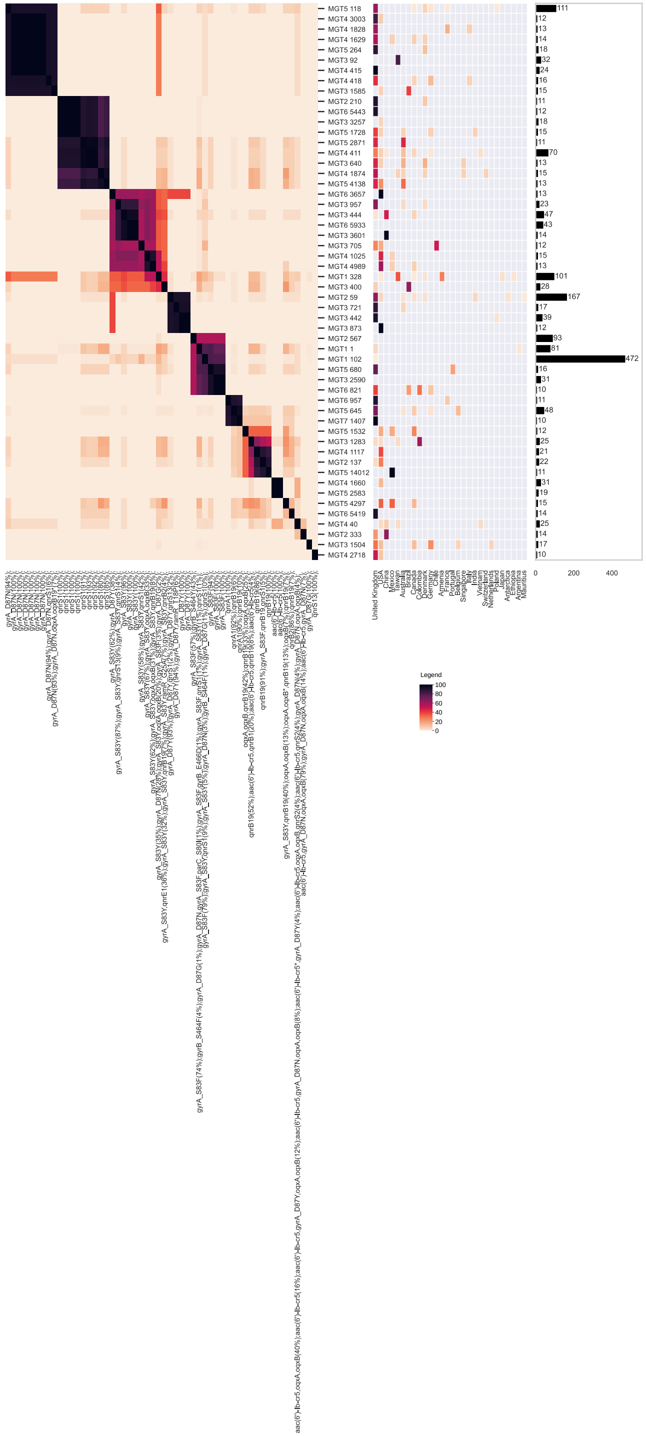

**Figure S9**.

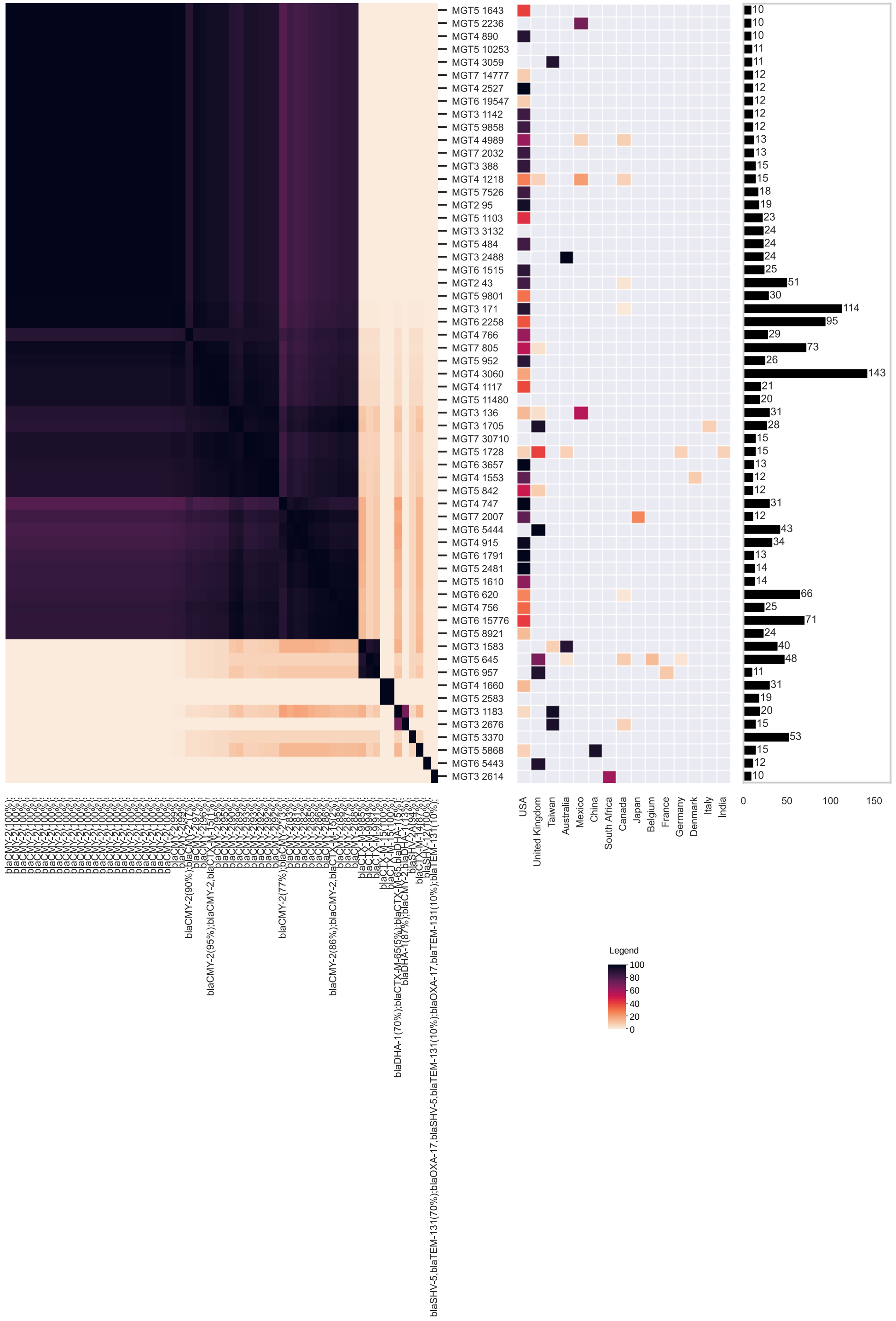
Legend
100
80
60
40
20
0
USA
United Kingdom
Taiwan
Australia
Mexico
China
South Africa
Canada
Japan
Belgium
France
Germany
Denmark
Italy
India
0
50
100
150

**Figure S10**.

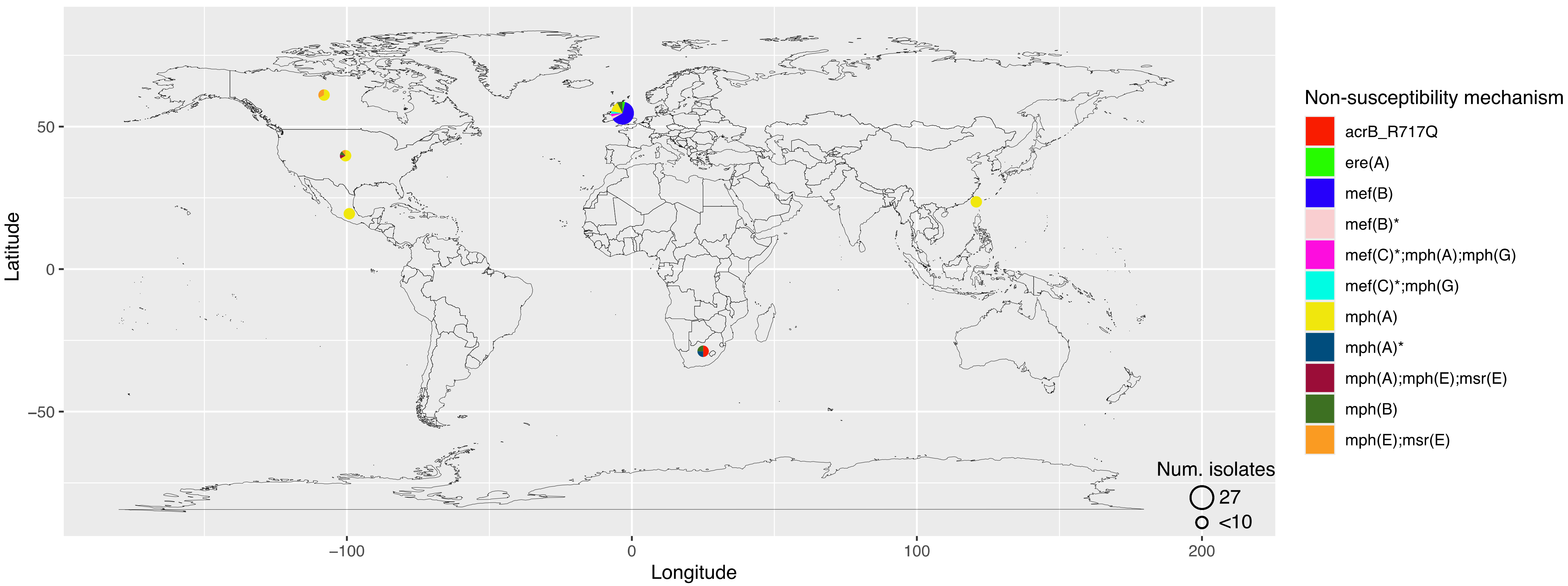
50
0
−50
Latitude
−100
0
100
200
Longitude
Num. isolates
27
<10
Non-susceptibility mechanism
acrB_R717Q
ere(A)
mef(B)
mef(B)*
mef(C)*;mph(A);mph(G)
mef(C)*;mph(G)
mph(A)
mph(A)*
mph(A);mph(E);msr(E)
mph(B)
mph(E);msr(E)

**Figure S11.**

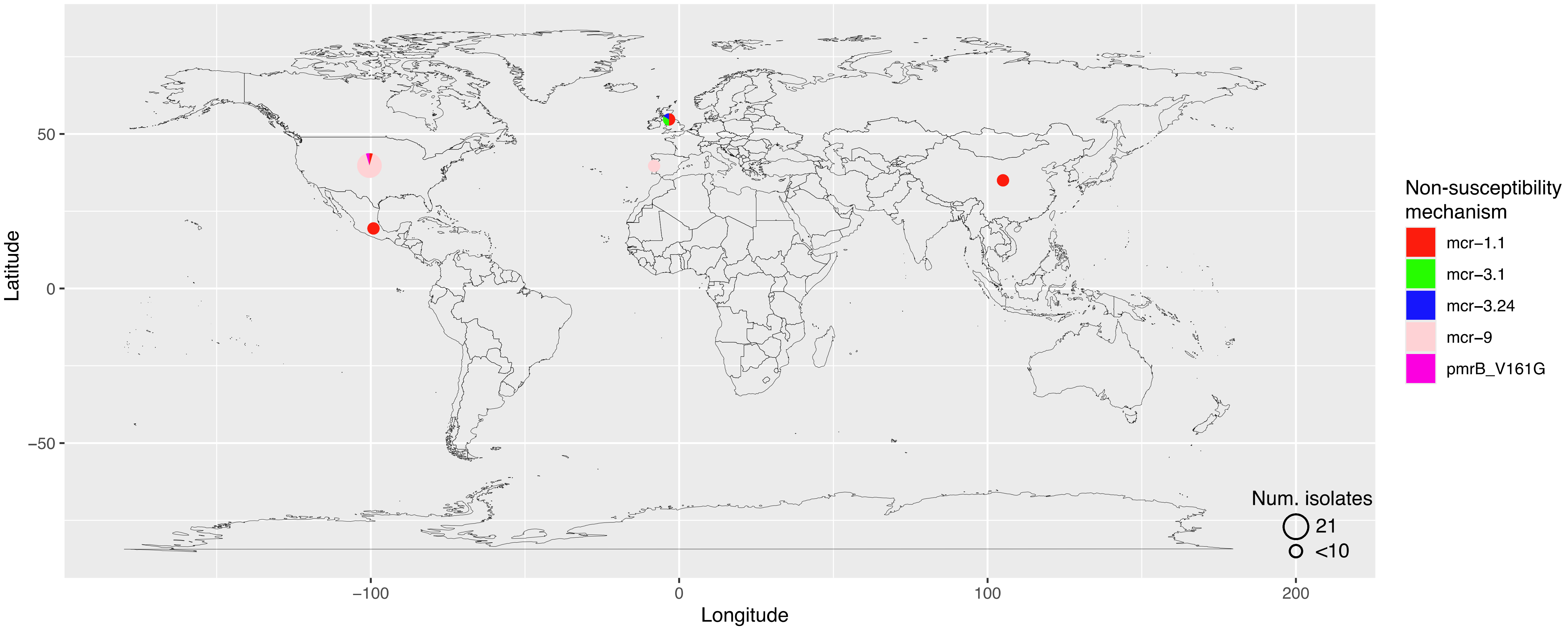

50
0
−50
Latitude
−100
0
100
200
Longitude
Non-susceptibility mechanism
mcr−1.1
mcr−3.1
mcr−3.24
mcr−9
pmrB_V161G
Num. isolates
21
<10


**Figure S12**.
